# Mechanistic dissection of the *CHD4* enhancers reveals cooperative functions among the homotypic ZNF410 clustered motifs

**DOI:** 10.1101/2024.02.24.581901

**Authors:** Siyuan Xu, Ren Ren, Chuxuan Peng, Haowen Lu, Lana C. Ly, Merlin Crossley, Bin Xu, Xiaodong Cheng, Gerd A. Blobel, Xianjiang Lan

**Author notes:** Correspondence (G.A.B.), (X.L.).

## Abstract

Transcription factors often regulate numerous target genes. However, ZNF410 controls only a single gene, *CHD4,* in human erythroid cells by its highly restricted chromatin occupancy to the *CHD4* locus via two clusters of ZNF410 binding motifs. Here, we uncover that ZNF410 controls chromatin accessibility and activity of the two *CHD4* enhancer regions. Combining CRISPR/Cas9 genomic deletion with CRISPRi approaches, we demonstrate that both enhancer regions additively contribute to *CHD4* gene expression. Mutations of the di-adenine nucleotides within the ZNF410 binding motif fully disrupt ZNF410-DNA interaction. Luciferase assays, ChIP-seq, and ATAC-seq studies reveal that the homotypic clustered motifs within the *CHD4* enhancers are recognized by ZNF410 in a collaborative fashion. Especially, the three ZNF410 motifs located in the 3’ end of the distal enhancer act as “switch motifs” to control the chromatin accessibility and activity of the whole distal enhancer region. Together, our findings expose a complex functional hierarchy of motifs, where clustered motifs bound by the same transcription factor cooperate to fine-tune the expression of target gene.

**Highlights:**

- ZNF410 maintains the accessibility of the *CHD4* enhancer regions
- The two enhancer elements additively modulate *CHD4* gene expression
- The dinucleotide “AA” is the key bases within ZNF410 binding motif
- ZNF410 binds to homotypic clustered motifs in a cooperative manner

## INTRODUCTION

The mammalian genome contains hundreds of thousands of *cis*-regulatory elements such as enhancers, silencers, and promoters, mediating the transcriptional outputs.^1^ Among these, enhancers are defined as *cis*-acting DNA sequences that can strengthen the transcription of target genes through binding of transcription factors (TFs) and thus increasing chromatin accessibility, regardless of the element’s location or orientation.^2^ Based on emerging technologies for characterizing enhancers at scale, candidate enhancers are identified using DNase I hypersensitivity (DHS),^3^ enhancer RNAs (eRNAs) transcription,^4^ enrichment of transcription co-activator p300,^5^ and chromatin histone modifications, such as H3K4me1 (histone H3 lysine 4 monomethylation)^6^ and H3K27ac (histone H3 lysine 27 acetylation).^7^ Despite that many studies have characterized genome-wide annotation of candidate enhancers,^5,6,8^ relatively few literatures shed insight into the mechanisms of the individual elements within these enhancers contributing to target gene expression in an additive,^9–11^ redundant,^12–14^ hierarchical,^15,16^ or synergistic manner^17–19^ in specific gene loci. Moreover, clustering of multiple transcription factor binding sites (TFBSs) for the same TF has proved to be a pervasive feature of enhancers/promoters in eukaryotic genome.^20^ However, the contribution of each binding site within the homotypic clusters of TFBSs (HCTs) to target gene expression in its native genomic context remains to be studied.

β-thalassemia and sickle-cell disease (SCD) are the genetic disorders with a worldwide distribution, caused by mutations in the human β-globin gene locus. Elevating fetal hemoglobin (HbF, consisting of two α and two γ chains) to replace adult hemoglobin (HbA, consisting of two α and two β chains) is an effective strategy for hemoglobinopathy treatments.^21,22^ Recently, multiple novel HbF repressors were identified via TF-based CRISPR/Cas9 genetic screens in HUDEP-2 cells, such as ZNF410,^23,24^ HIC2,^25^ and NFI.^26^ Among these, ZNF410 uniquely silences HbF through activating its sole target gene *CHD4*, a known HbF repressor in erythroid cells.^23,27^ Mechanistically, ZNF410 only binds to the *CHD4* regulatory regions harboring two highly conserved motif clusters in the genome, with each containing more than 10 copies of ZNF410 binding motifs, via its tandem array of five C2H2 zinc-finger (ZF) DNA binding domain.^23^ ZNF410 is a monomer and binds to motif-containing DNA oligonucleotide with 1:1 stoichiometry *in vitro*.^28^ However, the mechanisms of ZNF410 regulating *CHD4* gene expression through motif clusters are not understood.

Here, through chromatin immunoprecipitation with high-throughput sequencing (ChIP-seq), assay for transposase-accessible chromatin with high-throughput sequencing (ATAC-seq) analyses, in conjunction with loss-of-function experiments by CRISPR/Cas9 DNA fragment editing and CRISPR interference (CRISPRi), we show that ZNF410 maintains the accessibility of the *CHD4* regulatory regions composed of two enhancers, and both enhancers activate *CHD4* gene expression in an additive manner. Combining *in vitro* luciferase assays with episomal ChIP, in conjunction with ChIP-seq and ATAC-seq analyses using knock-in clones, we further demonstrate that ZNF410 binds to DNA via functional collaborations among the homotypic ZNF410 clustered motifs. Our findings uncover the mechanisms of ZNF410 binding to *cis*-regulatory elements to drive target gene expression, offering a paradigm regarding the singular motif specificity and motif orientation of HCTs recognized by TF.

## RESULTS

### ZNF410 generally regulates the *CHD4* gene expression

Recent studies have revealed that ZNF410 represses the fetal globin expression by singular activation of the nucleosome remodeling and deacetylase (NuRD) component *CHD4* through directly binding to the *CHD4* regulatory regions in human erythroid cells.^23,24^ However, it remains unclear whether ZNF410 regulating *CHD4* transcription is cell-type specific. Based on the analyses of Genotype-Tissue Expression (GTEx) dataset, ZNF410 may be generally limiting for *CHD4* gene expression.^23,24^ We also analyzed the correlation of *ZNF410* and *CHD4* according to the Human Protein Atlas (HPA) dataset,^29^ and again found the positive correlation of *ZNF410* and *CHD4*, but not *CHD3* (Figures S1A and S1B). We further examined the Tabula Sapiens dataset,^30^ a single-cell transcriptomic atlas of humans, and also observed high correlation of *ZNF410* and *CHD4* expression across the blood cells (Figures S1C). To confirm ubiquitous ZNF410-*CHD4* regulation, we performed ZNF410 ChIP-seq experiments and found that ZNF410 binds to the *CHD4* regulatory regions across all tested cell lines (Figure S1D). Importantly, depletion of ZNF410 significantly downregulated the *CHD4* gene expression in these cell lines (Figure S1E). These data together indicate that ZNF410 generally activates *CHD4* expression. To further explore the mechanisms of ZNF410-*CHD4* regulation, we focused on K562 cells, an erythroleukemia cell line.

### ZNF410 maintains the accessibility of the *CHD4* enhancer regions

To compare the binding pattern of ZNF410 between K562 and HUDEP-2 cells, we reanalyzed the ChIP-seq and ATAC-seq results of HUDEP-2 cells from previous studies.^23,26,31^ Next, we performed ChIP-seq experiments and found that the ZNF410 binding profile in K562 cells is similar to that of HUDEP-2 cells (Figure 1A and S1F). There are five ZNF410 peaks (hereinafter referred to as peak1 to peak5) in the *CHD4* regulatory regions. The similar ZNF410 binding profiles were detected and HOMER motif analysis based on the ChIP-seq data revealed that the derived ZNF410 binding motif is identical to that of HUDEP-2 cells (Figures 1A, S1F, and S1G).^23^ Of note, the binding intensity of peak4 is stronger than that of HUDEP-2 cells (Figure 1A). We also examined enrichment levels of active chromatin marks, H3K27ac and H3K4me3 (histone H3 lysine 4 trimethylation), by ChIP-seq and chromatin accessibility by ATAC-seq in K562 cells. We observed that similar to HUDEP-2 cells, the ZNF410 peaks fall into open chromatin and H3K27ac enriched regions (Figures 1A and S1F). In addition, the *CHD4* regulatory regions produce putative eRNAs and enrich H3K4me1, DNase I, EP300 and RNA polymerase II (Pol II) signals according to the cap analysis of gene expression (CAGE) and ENCODE databases (Figure S1H).^8,32^ These hallmarks of active enhancers indicate that the *CHD4* regulatory regions are composed of two enhancers, which we called the proximal enhancer and distal enhancer (Figures 1A and S1H).

**Figure 1.**
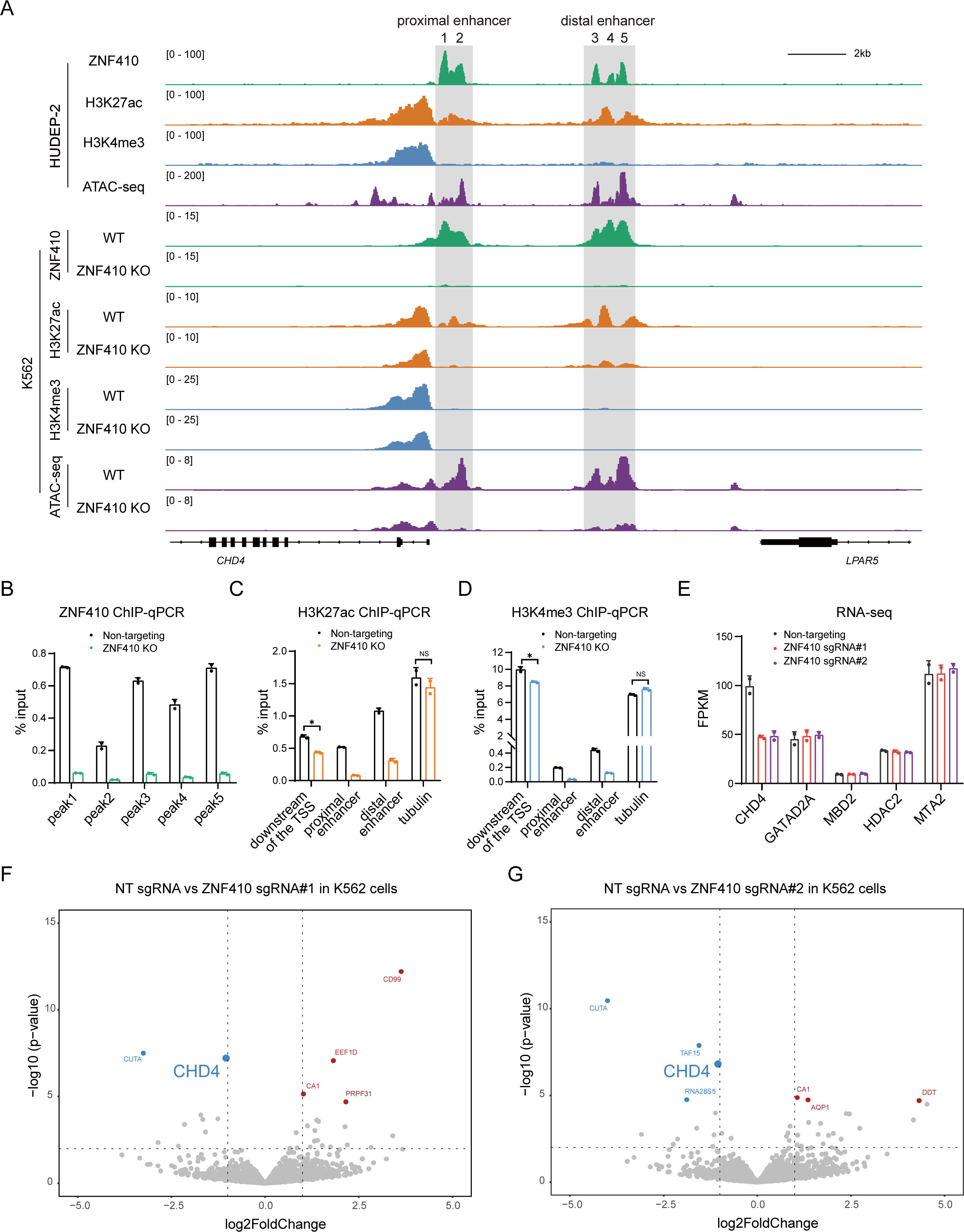
ZNF410 maintains the accessibility of the *CHD4* enhancer regions to drive gene expression. (A) ChIP-seq and ATAC-seq profiles in the *CHD4* enhancer regions in HUDEP-2 and K562 cells. Upper panel: shown are ZNF410 (green), H3K27ac (orange), H3K4me3 (light blue) ChIP-seq, and ATAC-seq (purple) in HUDEP-2 cells. Lower panel: anti-ZNF410, anti-H3K27ac, anti-H3K4me3 (light blue) ChIP-seq, and ATAC-seq tracks of wild-type (WT) and ZNF410-depleted K562 cells. KO, knockout. (B-D) ChIP-qPCR results of ZNF410, H3K27ac and H3K4me3 in the *CHD4* enhancer regions in WT and ZNF410-depleted K562 cells, respectively. Results are shown as mean ± SD (n=2). \**P* < 0.05, NS, Not Significant, unpaired Student’s *t*-test. (E) Expression levels of *CHD4*, *GATAD2A*, *MBD2*, *HDAC2*, *MTA2* by RNA-seq analysis of WT and ZNF410-depleted K562 cells (by ZNF410 sgRNA#1 and sgRNA#2). Results are shown as mean ± SD (n=2). FPKM, Fragments per kilo base per million mapped reads. (F-G) Volcano plot of RNA-seq analysis in WT and ZNF410-depleted K562 cells (by ZNF410 sgRNA#1 and sgRNA#2). NT, non-targeting, related to Figure S1.

We knocked out (KO) ZNF410 in K562 cells (Figure S1I) and carried out ChIP-seq and ChIP-qPCR experiments with a specific antibody against ZNF410, H3K27ac, and H3K4me3 respectively. We found that the depositions of H3K27ac and H3K4me3 are dramatically decreased in the proximal and distal enhancer regions in ZNF410-depleted K562 cells, whereas the H3K27ac and H3K4me3 levels had modest reduction at the downstream of the *CHD4* transcription start site (TSS) (Figures 1A-1D). Notably, the enrichment levels of H3K27ac and H3K4me3 had no changes in other ZNF410-binding loci after ZNF410 depletion (Figure S1F). Importantly, ATAC-seq results showed that the chromatin accessibility of the *CHD4* proximal and distal enhancer regions is nearly closed upon ZNF410 depletion, but no effect on the *CHD4* promoter (Figure 1A). Together, these findings imply that ZNF410 facilitates the accessibility of *CHD4* proximal and distal enhancers, whereas it is likely recruited to those open chromatin regions in the other six ZNF410-binding sites (*NBPF11*, *LIN54*, *TIMELESS*, *ICAM2*, *CBX8*, and *SUPT16H*) (Figure S1F). Next, we performed RNA-seq experiments to understand how ZNF410 regulates gene expression in ZNF410-depleted K562 cells. Notably, the expression of ZNF410 was significantly decreased and *CHD4* was the only ZNF410-regulated NuRD subunit (Figures 1E and S1J), which was further validated by RT-qPCR experiments (Figure S1K). According to differential gene expression analysis, with a threshold setting of at least 2-fold (*q* < 0.05) in both ZNF410 sgRNAs of the biological replicates (Figures 1F and 1G), we discovered that besides *CHD4*, there are two additional ZNF410-regulated genes *CA1* and *CUTA* (Table S1). However, no ZNF410 ChIP-seq signals were detected at their gene loci, indicating that they are not direct target genes. In addition, the transcriptional levels of other 6 ZNF410-bound genes by ChIP-seq were minimally affected (Figure S1L). Taken together, these findings uncover that ZNF410 maintains the accessibility of the *CHD4* enhancer regions to singularly activate *CHD4* gene expression in K562 cells.

### The *CHD4* enhancer elements act in an additive manner

Multiple enhancers for a single gene can additively, redundantly, or cooperatively function to influence the gene expression.^1,33–35^ Previous report has found that deletion of a 6.7 kb DNA fragment containing both *CHD4* enhancers leads to similar reduction of *CHD4* transcription levels compared to that by ZNF410 KO in HUDEP-2 cells.^24^ To investigate the individual contribution of the proximal and distal enhancers in regulating *CHD4* gene expression, we first specifically deleted the ZNF410 binding regions of each enhancer by CRISPR DNA fragment editing in K562 cells (Figure 2A). Given that *Chd4* knockout blastocysts do not implant and are hence not viable,^36^ we designed sequence-specific single-guide RNAs (sgRNAs) deleting the partial core promoter downstream of the *CHD4* TSS as a positive control to compare with the effect of individual enhancer deletion (Figures 2A, 2B, S2A, and S2E). We isolated three independent homozygous single-cell CRISPR deletion clones (Figures S2B, S2C, S2F, and S2G). RT-qPCR analysis of these single-cell clones revealed that deletion of the proximal enhancer results in approximately 32% reduction of *CHD4* mRNA levels and loss of the distal enhancer leads to a reduction of 20% (Figures 2C and 2D).

**Figure 2.**
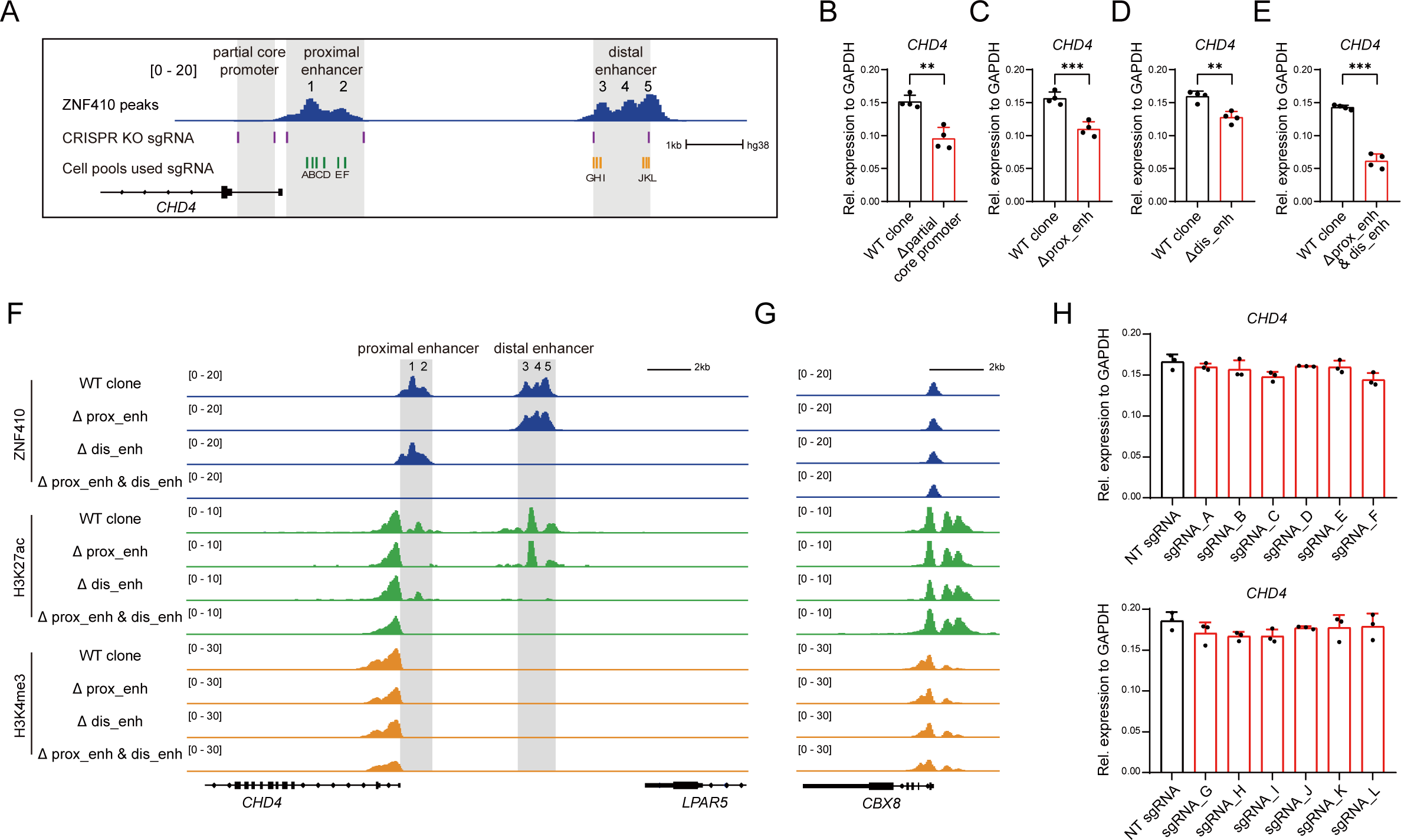
CRISPR genetic and epigenetic perturbation of the *CHD4* proximal and distal enhancers. (A) Schematic of the sgRNAs location in the *CHD4* proximal and distal enhancers used by CRISPR DNA fragment editing and CRISPRi perturbation. (B-E) Expression levels of *CHD4* by RT-qPCR analysis of WT (n=4) and single- and double-enhancer knockouts clones (n=4). (B) partial core promoter deletion clones, (C) single proximal enhancer (prox_enh) deletion clones, (D) single distal enhancer (dis_enh) deletion clones and (E) double enhancer (prox_enh & dis_enh) deletion clones, respectively. Results are shown as mean ± SD. \*\**P* < 0.01, \*\*\**P* < 0.001; unpaired Student’s *t*-test. (F-G) ChIP-seq tracks of ZNF410 (dark blue), H3K27ac (green), and H3K4me3 (orange) of WT and single- and double-enhancer knockouts clones in the *CHD4* proximal and distal enhancers (F) and the *CBX8* locus (G). (H) RT-qPCR analysis of *CHD4* mRNA levels of K562 Cas9 stable cell line pools transduced with the indicated sgRNAs. Results are shown as mean ± SD (n=3), related to Figure S2.

To further discriminate the possibility of the proximal and distal enhancers regulating *CHD4* expression in an additive, redundant, or synergistic manner, we constructed proximal and distal enhancers double knockout (DKO) subclones using the distal enhancer deletion clone (Figures S2D and S2H). The *CHD4* mRNA levels were decreased approximately 57% compared with wide-type (WT) clones, suggesting that the proximal and distal enhancers regulate *CHD4* in an additive manner (Figure 2E). The reduced *CHD4* mRNA levels were comparable to that with ZNF410 depletion (Figures 1E and S1K). Notably, the expression levels of other NuRD complex subunits (*MBD2* and *HDAC2*) were not affected in all cases with the DNA element deletions (Figures S2I-S2L). We next performed ChIP-seq experiments with anti-ZNF410 and found that the binding regions of ZNF410 is successfully deleted by CRISPR mediated genomic engineering (Figure 2F). Meanwhile, the active enhancer marker H3K27ac signals were correspondingly abolished in the proximal enhancer deletion clones, distal enhancer deletion clones, and DKO subclones, respectively (Figure 2F). We also found that deletion of the enhancer elements results in a reduction of H3K4me3 occupancy at the *CHD4* promoter regions (Figure 2F). Of note, no off-target effect was observed in the *CBX8* locus (Figure 2G).

CRISPR genetic deletion perturbs the DNA sequences of the enhancer regions. As an alternative to genetic deletion, we leveraged a CRISPRi approach to silence the *CHD4* regulatory regions *in situ*.^37^ Given that there exists a 5’-CCC-3’ protospacer adjacent motif (PAM) in ZNF410 motif (Figure S1G), and then the dCas9-KRAB fusion protein would be recruited by the sgRNAs to block the ZNF410 binding (Figure 2A). To this end, we chose 12 sgRNAs locating across the center of ZNF410 binding peaks according to the ChIP-seq signals (six sgRNAs for the proximal enhancer and six sgRNAs for the distal enhancer, each sgRNA targets one motif, respectively) and transduced the indicated sgRNAs in K562 cell line stably expressing dCas9-KRAB and obtained positive cell populations by GFP or mCherry-based fluorescence-activated cell sorting (FACS) (Figure 2A). As expected, the *CHD4* mRNA levels were reduced when using the indicated sgRNAs to block the ZNF410 binding via CRISPRi (Figures S2M and S2N). The expression levels of other NuRD complex subunit (*MBD2* and *HDAC2*) were not affected by CRISPRi (Figures S2M and S2N), implying specific perturbations. We noted that the effect of sgRNAs targeting the proximal enhancer on *CHD4* gene expression is stronger than that of the distal enhancer, consistent with the results from enhancer deletion clones (Figures 2C and 2D). We next carried out ZNF410 ChIP-seq experiments and found that blocking the ZNF410 binding with sgRNAs targeting the proximal enhancer has no effect on that of its binding in the distal enhancer, and vice versa (Figures S2O and S2P). Notably, recruiting dCas9-KRAB to peak1 of the proximal enhancer can partially interrupt ZNF410 binding in both peak1 and peak2 regions, and targeting the peak5 of the distal enhancer can impair ZNF410 binding in all peak3, peak4, and peak5 regions (Figures S2O and S2P) (see below).

Based on these observations, we next utilized pairs of sgRNAs simultaneously targeting the proximal and distal enhancers and found a greater decrease of *CHD4* mRNA levels (Figure S2Q), consistent with the results from ZNF410 deficient cells and DKO subclones (Figures 1E, S1K, and 2E). Again, the expression levels of *MBD2* and *HDAC2* were not affected (Figure S2Q). We further performed ChIP-seq experiments with ZNF410 antibody and found that the enrichment of ZNF410 is apparently decreased in the proximal and distal enhancer (Figure S2R). Importantly, the deposition of active enhancer marker H3K27ac was nearly abolished in these regions (Figure S2R). In contrast, the repressive histone mark H3K9me3 (histone H3 lysine 9 trimethylation) signals were significantly established across the enhancer regions (Figure S2R), due to recruitment of KAP1 and SETDB1.^38,39^ We also observed that the enrichment of H3K4me3 was modestly decreased at the *CHD4* promoter region (Figure S2R).

While blocking the ZNF410 binding via CRISPRi approach, the adjacent TF bindings would be also interrupted. Therefore, we further employed CRISPR/Cas9 to disrupt a single ZNF410 binding site to explore whether destructing a single ZNF410 binding motif in the proximal or distal enhancer would affect *CHD4* gene expression. To this end, we transduced the indicated sgRNAs into K562 cells stably expressing Cas9 (Figure 2A) and then measured the editing efficiency of each sgRNA with 70%-90% indels (Figure S2S). Of note, each sgRNA would cause a cleavage between the di-adenine nucleotides within the ZNF410 binding motif, further producing indels to abolish ZNF410 binding (see below). Compared with blocking of the ZNF410 binding in the proximal or distal enhancers, impairment of any single ZNF410 binding motif had minimal effect on *CHD4* mRNA levels when using the same sgRNAs (Figure 2H), as well as *MBD2* and *HDAC2* mRNA levels (Figure S2T). Collectively, these data indicate that the proximal enhancer and distal enhancer contribute to the *CHD4* gene expression in an additive manner. Moreover, interference of the clustered ZNF410 motifs in the proximal and/or distal enhancers, but not a single one, diminishes *CHD4* gene expression.

### Perturbation of the *CHD4* enhancer regions induces g-globin levels

We previously reported that ZNF410 depletion elevates γ-globin levels by modulating *CHD4* expression in erythroid cells.^23^ Since several studies including ours have found that the basal levels of γ-globin levels in HUDEP-2 single-cell clones vary dramatically,^26,40^ thus we employed HUDEP-2 cell pools to explore the sensitivity of γ-globin genes to *CHD4* levels and validate the results of K562 cells through manipulating the ZNF410 binding to the *CHD4* enhancers. Firstly, indicated sgRNAs were individually transduced into HUDEP-2 cell line stably expressing Cas9 to disrupt a single ZNF410 binding motif in the proximal enhancer or distal enhancer. RT-qPCR experiments revealed that the mRNA levels of *CHD4* and γ-globin had minimal change (Figures S3A and S3B). We also examined the α-globin and *GATA1* mRNA levels, and found no change (Figures S3A and S3B), indicating that disruption of individual ZNF410 binding motifs does not impair erythroid differentiation, consistent with our previous findings.^23^ Secondly, we individually transduced the same sgRNAs into HUDEP-2 cell line stably expressing dCas9-KRAB to block ZNF410 binding in the proximal enhancer or distal enhancer. As expected, we found a greater decrease of *CHD4* mRNA levels and correspondingly stronger increase of γ-globin levels by RT-qPCR experiments (Figures S3C and S3D), compared to the results of a single ZNF410 binding motif disruption by Cas9. Again, the mRNA levels of α-globin and *GATA1* were unaffected (Figures S3C and S3D). The *CHD4* enhancer regions activated gene expression in an additive manner (Figure 2). Therefore, simultaneously silencing the proximal and distal enhancers would cause an additive or synergistic increase of γ-globin mRNA levels. Finally, we utilized pairs of sgRNAs concurrently targeting the proximal and distal enhancers to block the ZNF410 binding, which caused approximately 60% decrease of *CHD4* mRNA and thus a robust increase of γ-globin mRNA (Figure S3E), similar to the results in ZNF410-deficient HUDEP-2 cells.^23^ Of note, *CHD4* dosage changes had overall nonlinear effect on γ-globin gene expression. Importantly, silencing the *CHD4* enhancer regions did not affect erythroid maturation (Figure S3E). Consistent with the results by RT-qPCR, western blotting revealed that the γ-globin protein levels are also significantly elevated without affecting the ZNF410 protein levels (Figure S3F). In sum, these data indicate that perturbation of the *CHD4* enhancer regions elevate γ-globin gene expression in a *CHD4* dosage-dependent manner.

### Characterization of conserved homotypic ZNF410 clustered motifs at the *CHD4* **enhancers in mammals**

ZNF410 orthologs are highly conserved in mammals, suggesting that ZNF410 likely retains the function of binding to the *CHD4* enhancers in the evolution.^28^ Our ChIP-seq analysis revealed that the *CHD4* enhancers contain five ZNF410 peaks (Figure 1A), harboring 22 ZNF410 motifs (Figure 3A).^23,24^ Of note, additional 5 motifs are outside the ZNF410 peaks (Figure 3A). These clustered ZNF410 motifs are composed of two HCTs within the proximal enhancer and distal enhancer respectively. We examined the amount of ZNF410 motifs in each peak and the distance between ZNF410 motifs in human (Figures 3A and S4A). Notably, each peak covers at least three ZNF410 motifs, which we defined as a sub-HCTs. We also examined the homotypic Zfp410 clustered motifs at the *Chd4* locus in mouse and found that the amount of Zfp410 motifs and the distance between Zfp410 motifs are highly similar compared to those in human (Figures 3A and S4A). Of note, unlike the *CHD4* enhancers, each of the other 6 ZNF410 binding regions (*NBPF11*, *LIN54*, *TIMELESS*, *ICAM2*, *CBX8*, and *SUPT16H*) only contains one binding motif (Figure S1F). Thus, ZNF410 maintaining the chromatin accessibility to activate gene expression appeared to rely on the homotypic clustered motifs, but not a single one (Figures 3A and S1F). Moreover, we analyzed the homotypic ZNF410 orthologs clustered motifs at the *CHD4* locus in the genomes of other mammals including pig and dog, and found that the ZNF410 binding motifs are also highly conserved, especially the HCTs in the distal enhancer (Figures 3A and S4A). Multiple sequence alignment of the *CHD4* enhancers revealed the similarity of ZNF410 binding regions (Figure S4B). Notably, neither the number of ZNF410 motifs nor the distance between motifs is conserved in non-mammalian vertebrates such as zebra finch in Aves, African clawed frogs in Amphibia, and zebrafish in Osteichthyes (Figures S4C and S4D).

**Figure 3.**
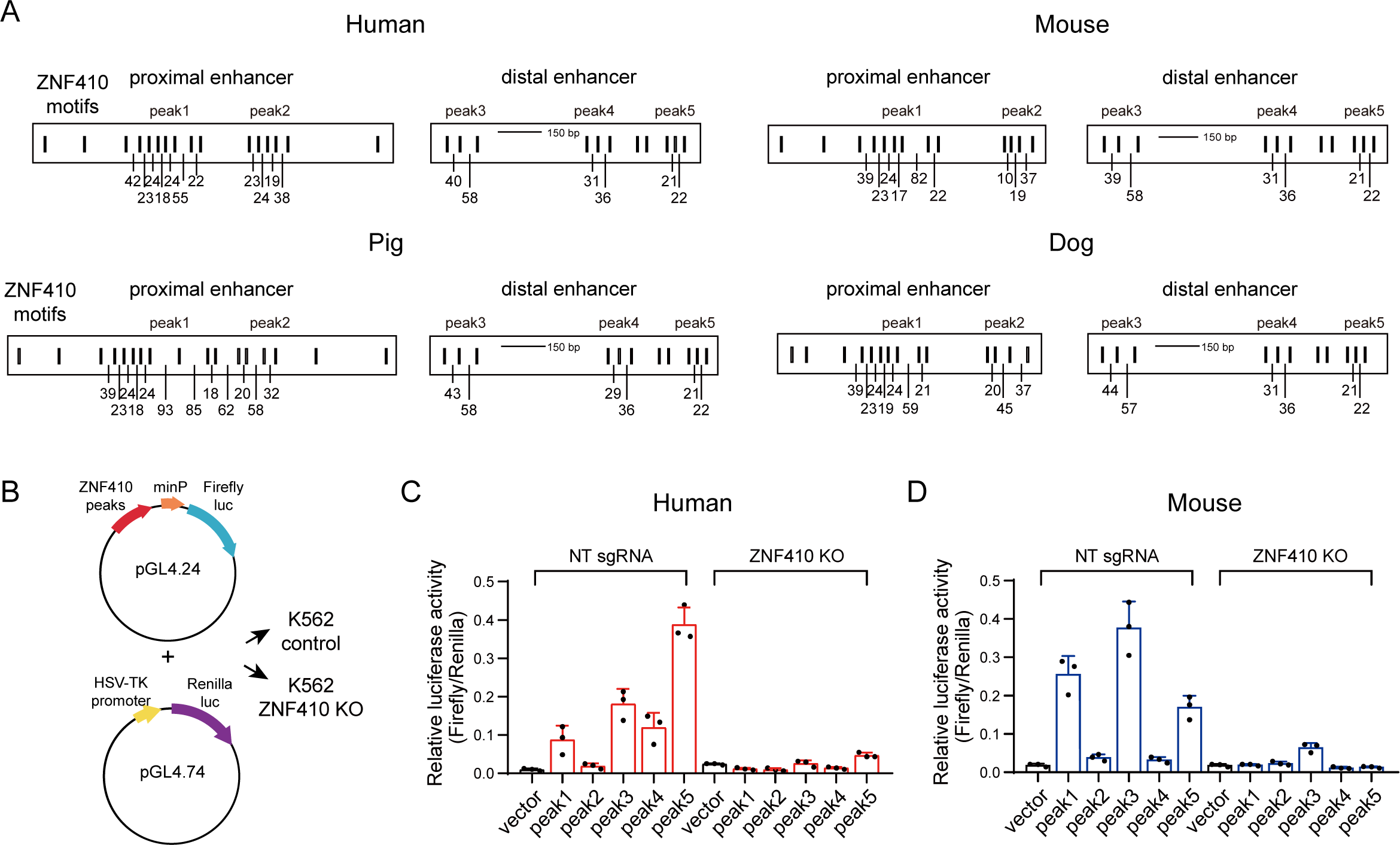
Conservation and function of the homotypic ZNF410 clustered motifs. (A) The pattern of ZNF410 binding motifs in the human, mouse, pig, and dog *CHD4* enhancer regions. The numbers indicate the distances of adjacent ZNF410 motifs. (B) Schematic of luciferase assays of WT and ZNF410 KO K562 cells. Each ZNF410 peak was cloned into pGL4.24 plasmid. Renilla luciferase activity was used as a normalization for transfection efficiency. (C) Results of luciferase reporter assay to measure each human ZNF410 peak activity in WT and ZNF410 KO K562 cells. Results are shown as mean ± SD (n=3). NT, non-targeting. (D) Results of luciferase reporter assay to measure each mouse ZNF410 peak activity. Results are shown as mean ± SD (n=3), related to Figure S4.

To test whether the sub-HCTs within each peak are capable to drive transcription, the five ZNF410 peaks of human and mouse were individually cloned into a luciferase reporter plasmid with a mini-promoter that enables basal gene expression. We next transfected these vectors into WT or ZNF410 KO K562 cells respectively (Figure 3B). Luciferase activity measurements revealed that each peak can more or less activate the reporter gene expression in a ZNF410-dependent manner (Figures 3C and 3D). Of note, the human peak2 and the mouse peak2 and peak4 displayed little ability to activate luciferase expression (Figures 3C and 3D), consistent with their weak ZNF410 binding intensity *in vivo* by the ChIP-seq results (Figures 1A and S4E).^23^ Moreover, the relative luciferase activity of the human peaks showed positive correlation with the sequence conservation (Figure S4F). Taken together, these data indicate that the ZNF410 HCTs within the *CHD4* enhancers are highly conserved in mammals and important contributors to the function of *cis*-regulatory sequences to promote gene expression.

### Interrogation of the key nucleotides within the motif for ZNF410 binding to DNA

We previously demonstrated that ZNF410 can directly bind to DNA with its tandem ZF domain by the crystal structure.^23^ However, the key nucleotides of ZNF410 motif for DNA binding need to be further validated. To this end, we preformed electrophoretic mobility shift assay (EMSA) with nuclear extracts from COS-7 cells overexpressing FLAG-tagged ZNF410 full-length (FL) or five tandem ZFs (ZF1-5), and found that the ZNF410-DNA binding is nearly disrupted when “AAT” nucleotides at the position from 10^th^ to 12^th^ of the motif were mutated to “GGG” (Figure S5A). Moreover, we mutated a single nucleotide each time to further gain insight into ZNF410-DNA binding. Importantly, point mutation of A10G or A11G, rather than A13G, in the ZNF410 motif completely blocked the binding to DNA (Figures 4A and 4B). We further confirmed the findings by more quantitative fluorescence polarization assays with the condition of 300 mM NaCl (Figures 4C and 4D). According to our structural data, the dinucleotide “AA” within ZNF410 motif corresponds to ZF2 that is essential for DNA binding (Figure S5B).^23^ Interestingly, we tested a natural A10C variant existed in each of the homotypic ZNF410 clustered motifs at the *GALNT18* locus that is the only locus containing ZNF410 HCTs besides the *CHD4* regions genome-wide,^23,24^ and found that this point mutation nearly lowers the luciferase activity to basal levels (Figures S5C and S5D), which we speculate is likely the reason why there is no ZNF410 binding at this site. In addition, we performed episomal ZNF410 ChIP-qPCR assays^41^ to further assess the binding in K562 cells transfected with luciferase vectors containing WT or MT ZNF410 motifs and found that replacing “AA” with “GG” within the motif significantly diminishes ZNF410 binding (Figures 4E and 4F), consistent with the results *in vitro*. Together these results reveal that the dinucleotide “AA” is the key bases mediating ZNF410 binding.

**Figure 4.**
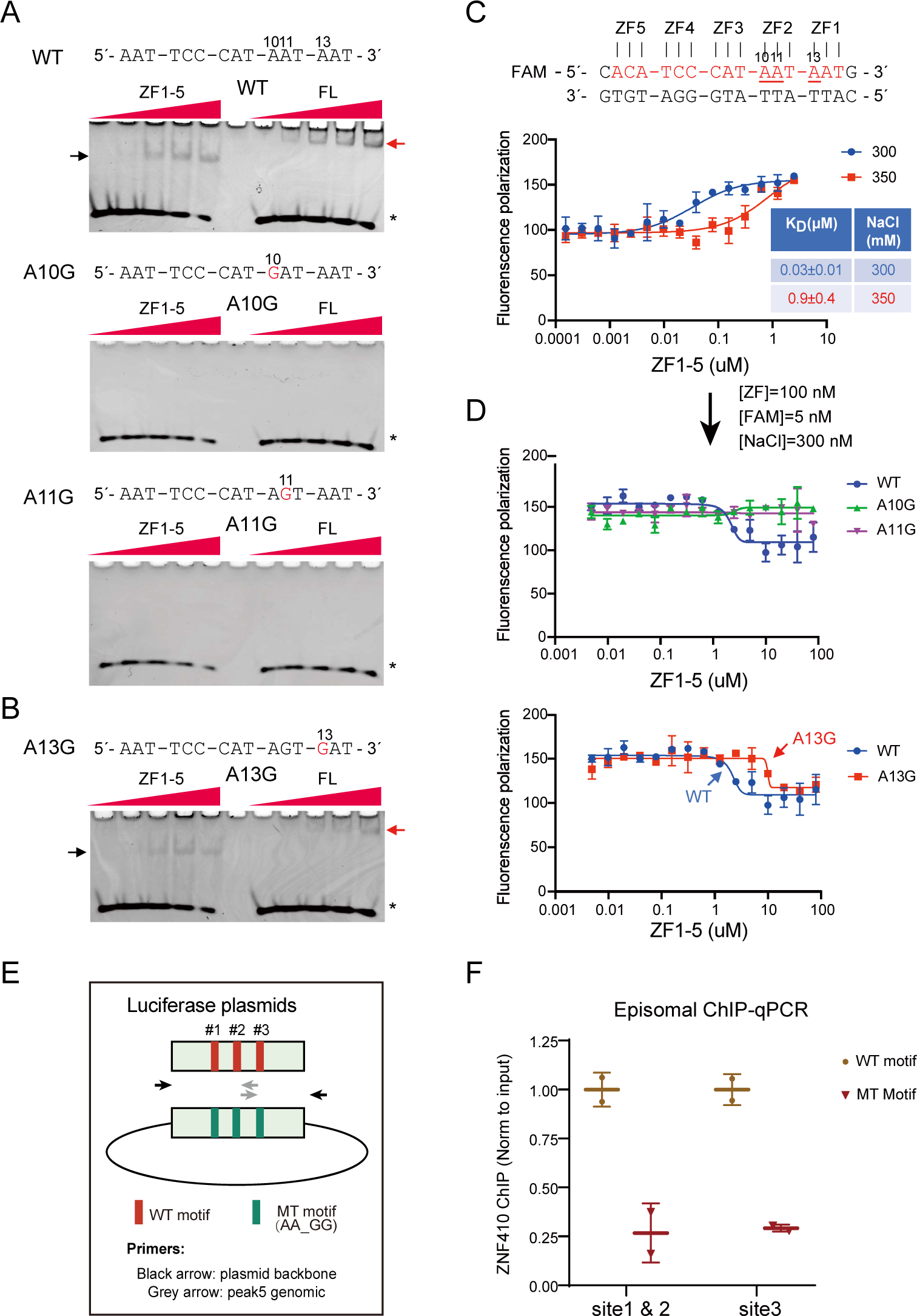
Identifying the key nucleotides of ZNF410 binding motif. (A-B) Determination of the key nucleotide of ZNF410 binding motif via EMSA experiments. EMSA probe sequences are shown above and mutant nucleotides are highlighted in red. The numbers are corresponding to nucleotide positions in the ZNF410 motif. Black arrow, ZF-probe complex; red arrow, ZNF410-probe complex; *, free probes; ZF, zinc finger; FL, full-length. (C) Binding affinity measurements of the ZF-DNA interaction by fluorescence polarization assay. Results are shown as mean ± SD (n=3). (D) Fluorescence polarization assay of the dissociation constants (K_D_) of the point mutation of A10G, A11G, or A13G within ZNF410 motif. Results are shown as mean ± SD (n=3). (E) Schematic of plasmids and primers for episomal ChIP-qPCR for WT and ZNF410 mutant motif plasmids. (F) Episomal ZNF410 ChIP-qPCR for K562 cells transfected with the plasmids in (E). Results are shown as mean ± SD (n=2), related to Figure S5.

### ZNF410 sub-HCTs exhibit cooperative binding *in vivo*

HCTs are a pervasive feature of human promoters and enhancers.^20^ Highly clustered motifs may promote cooperative recruitment of TFs, thus resulting in an all-or-none occupancy. As opposed to this scenario, each TF binding site independently contributes to gene expression.^42^ However, there are limited literatures studying the contribution of each motif within the HCTs to gene expression in cells. The human peak5, harboring ZNF410 sub-HCTs that displays robust luciferase activity, was considered as an ideal model (Figure 3C) to explore the function of each motif and also discriminate the two possibilities (all-or-none or independent occupancy) for ZNF410 binding. To this end, we constructed a series of luciferase reporter plasmids containing various combinations of mutated ZNF410 motifs by replacing “AA” with “GG” to investigate the effect of each ZNF410 motif on transcription (Figure 5A). Compared to WT peak5, mutating all three motifs (MT1) completely lost its ability to enhance the luciferase activity, indicating that ZNF410 binding is essential to drive transcription. Mutating any two motifs also displayed little activity, similar to empty vector (Figure 5A), implying that a single functional ZNF410 motif is insufficient to activate gene expression, in line with the cases of endogenous target genes (Figures S1F and S1L). Surprisingly, 3’-ZNF410-motif mutation (MT4) alone was sufficient to abolish the luciferase activity while mutating the 5’ motif (MT2) or middle motif (MT3) diminished the activity to some extent (Figure 5A), exhibiting a site-dependent function. Indeed, we observed the same phenomenon for both peak3 and peak4 (Figure S6A).

**Figure 5.**
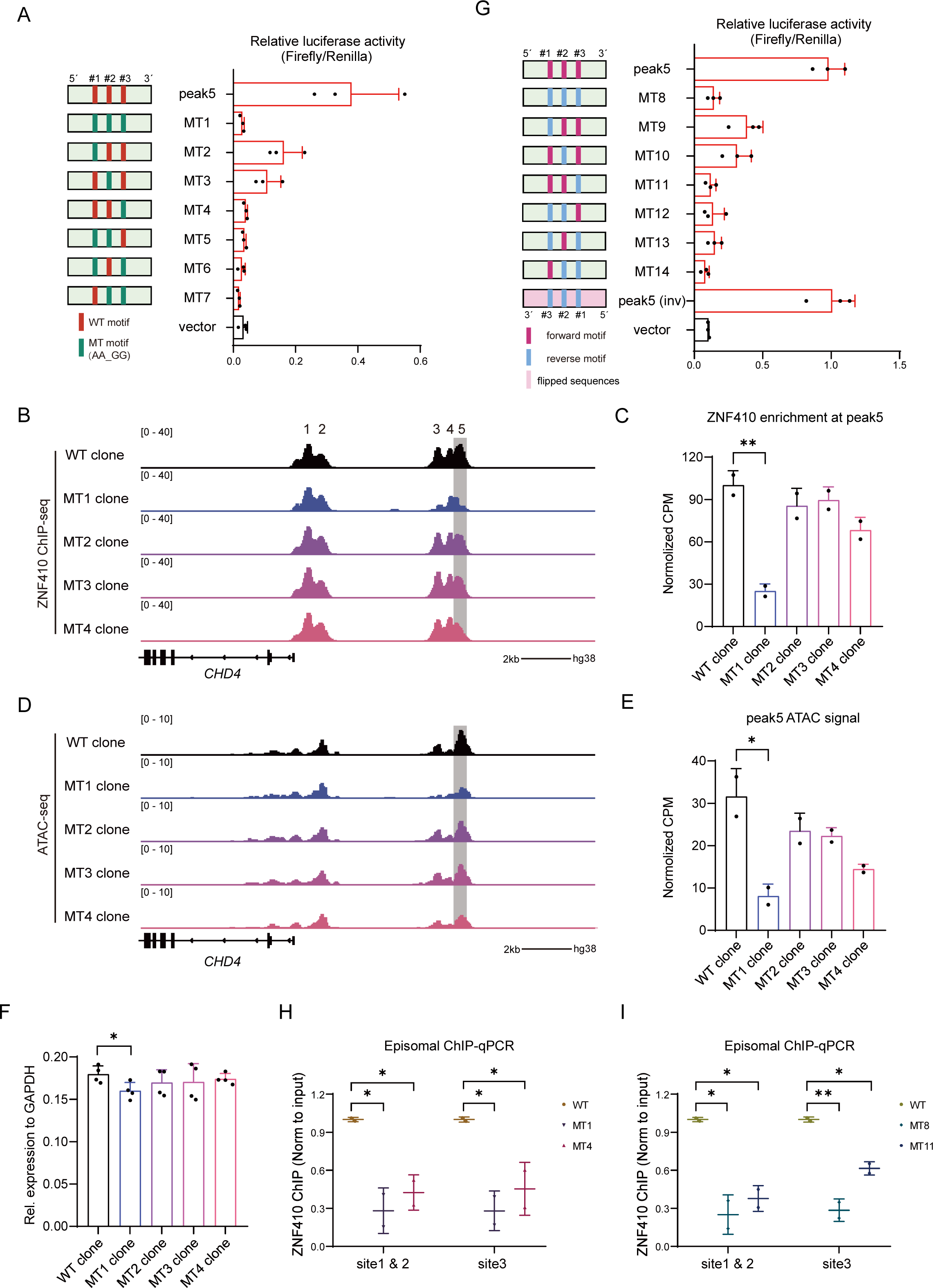
ZNF410 binds to sub-HCTs in a cooperative manner. (A) Schematic of all the variants of ZNF410 motif mutant sequences and results of luciferase reporter assays to measure the activity of all the variants of ZNF410 motif mutant sequences (n=3). MT, mutation. Results are shown as mean ± SD (n=3). (B) ChIP-Seq tracks of ZNF410 occupancy in the *CHD4* enhancer regions of each MT clones. (C) Bar plot depicts the mean ChIP-seq signal over peak5. Results are shown as mean ± SD (n=2). \*\**P* < 0.01, unpaired Student’s *t*-test. (D) ATAC-seq profiles in the *CHD4* enhancer regions of each MT clones. (E) Bar plot depicts the mean ATAC-seq signal over peak5. Results are shown as mean ± SD (n=2). \**P* < 0.05, unpaired Student’s *t*-test. (F) Expression levels of ZNF410 motif mutation homozygous single-cell clones. Results are shown as mean ± SD (n=4). \**P* < 0.05, unpaired Student’s *t*-test. (G) Schematic of the changes of ZNF410 binding motifs orientation sequences and results of luciferase reporter assays to measure the activity of the changes of ZNF410 binding motifs orientation sequences. Results are shown as mean ± SD (n=3). inv, inversion. (H) Episomal ZNF410 ChIP-qPCR for K562 cells transfected with the WT, MT1, and MT4 luciferase plasmids. Results are shown as mean ± SD (n=2). \**P* < 0.05, unpaired Student’s *t*-test. (I) Episomal ZNF410 ChIP-qPCR for K562 cells transfected with the WT, MT8, and MT11 luciferase plasmids. Results are shown as mean ± SD (n=2). \**P* < 0.05, \*\**P* < 0.01, unpaired Student’s *t*-test, related to Figure S6.

To further confirm our observations *in vivo*, we isolated two single-cell clones containing indels in the 3’ ZNF410 motif of peak5 (Figure S6B) and examined the ZNF410 enrichment by ChIP-qPCR experiments. As expected, 3’-ZNF410-motif impairment significantly decreased ZNF410 binding in peak5, whereas it had no effect on peak1 to peak4 in these clones (Figure S6C). ln the meantime, we knocked in “GG” over “AA” in the ZNF410 motifs of peak5 by CRISPR-mediated homology-directed repair and obtained four types of clones (two clones/type) containing either single or three ZNF410 mutated motif(s) (Figure S6D). ChIP-seq and ChIP-qPCR results revealed that the 3’-ZNF410-motif mutation (MT4 clone) results in a marked reduction of ZNF410 enrichment in peak5 while 5’-ZNF410-motif (MT2 clone) or middle-motif mutation (MT3 clone) had minimal effect, confirming that ZNF410 motifs exhibit a site-dependent role on binding to DNA (Figures 5B, 5C, S6E, and S6F), consist with the results of luciferase assay (Figure 5A). Of note, the WT ZNF410 binding motifs surrounding the mutant one might be also enriched by ZNF410 antibody after genomic DNA was randomly fragmented by sonication, resulting in residual ZNF410 ChIP-seq signals at peak5 in MT4 clone (Figures 5B and 5C). Moreover, we performed ATAC-seq experiments with these clones and found that the accessibility of peak5 is strongly reduced in MT4, but not MT2 or MT3 clone (Figure 5D and 5E). Of note, no effect was detected for peak1, peak2, peak3 and peak4 in these clones. More importantly, the ZNF410 enrichment levels and the accessibility of peak3, peak4, and peak5 were strikingly decreased upon mutating all three ZNF410 motifs in peak5 (MT1 clone) (see below). Lastly, RT-qPCR analysis with these single-cell clones revealed that mutating all three ZNF410 motifs in the peak5 (MT1 vs WT clone) leads to significant reduction of *CHD4* mRNA levels (Figure 5F), despite to a lesser degree than that of the distal enhancer deletion clones (Figure 2D).

Intriguingly, the ZNF410 HCTs at the *CHD4* locus have the same orientation, we next sought to explore the orientation effect of the ZNF410 motifs on transcription in the context of the peak5. To this end, we performed luciferase assays using a series of luciferase reporter plasmids containing forward or reverse ZNF410 motifs and found that the changes of ZNF410 motif orientation affect luciferase activity to varying degrees (Figure 5G). Notably, reversing the orientation of all three ZNF410 motifs completely abolished the luciferase activity (Figure 5G). We also observed the site-dependent effect (MT9 vs MT10 vs MT11) on the luciferase activity upon the changes of single ZNF410 motif orientation (Figures 5G and S6G), similar to those of mutating corresponding ZNF410 motif (Figures 5A and S6G). However, compared with peak5 or 3’-ZNF410-motif mutation (MT4), inversion of the whole sequences (peak5 vs peak5 inv or MT4 vs MT4 inv) had no effect on luciferase activity (Figures 5G and S6H), in line with that enhancer in the context of plasmid stimulates gene expression in an orientation-independent manner.^2^ Together these data show that ZNF410 motif orientation has a significant effect on gene transcription, likely due to alteration of flanking DNA that is critical for TF binding.^43,44^

Based on these results, we speculated that ZNF410 binding to the 3’ motif of peak5 assists its binding to the other motifs (Figure 5A), further conferring site-dependent role. To verify our hypothesis, we first carried out EMSA experiments with 120 bp free DNA probes containing all three ZNF410 motifs (core sequence of peak5) (Figures S6I-S6L). As expected, both purified ZF1-5 and FL bound to the WT probe, resulting in three types of migrating protein-DNA complexes as the concentration increases by native gel electrophoresis, respectively (Figures S6K and S6L). We next incubated ZF1-5 or FL with DNA probes mutating 5’, middle or 3’ motif, and found that all these mutant probes formed two types of protein-DNA complexes in a similar pattern, indicating that disrupting any single motif failed to impair the binding to the other two functional motifs (Figures S6K and S6L). Prior studies have indicated that nucleosome-mediated cooperation of TFs binding to DNA could not be detected *in vitro* using naked DNA templates.^45–48^ To determine the effect of nucleosomes on ZNF410–DNA binding, we reconstituted a 200 bp DNA sequence covering the three ZNF410 motifs of peak5 into a nucleosome using purified recombinant human histones (Figures S6J and S6M). The 200 bp DNA region was enriched with ZNF410 binding signals and accessible according to ZNF410 ChIP-seq and ATAC-seq data (Figure S6J), and it can accommodate only one nucleosome containing both nucleosomal and free DNA with ZNF410 motifs (Figure S6M). Of note, reconstituted nucleosome with 3’ free DNA shifts slower than that with 5’ free DNA.^49^ Notably, EMSA assays detected only one protein-nucleosome complex, indicating that ZF1-5 can only bind to the 3’ free DNA with motif (Figure S6M). These observations argue against ZNF410 as a pioneer factor,^50^ consistent with a previous study by nucleosome consecutive affinity purification–systematic evolution of ligands by exponential enrichment (NCAP– SELEX) approach.^51^ These data also indicate that ZF1-5 binding to 3’ free DNA is unable to promote its binding to the other two motifs on nucleosomal DNA *in vitro*, and other unknown co-factors are likely required for this cooperation.

To test the hypothesis *in vivo*, we again utilized episomal ChIP-qPCR assays in K562 cells transfected with luciferase vectors containing WT peak5, MT1, or MT4. As expected, mutating all motifs (MT1) strongly reduced ZNF410 binding to these sites, compared with WT. Importantly, not only impaired the binding to itself (site 3), 3’-motif mutation (MT4) also lead to significant reduction of the ZNF410 binding to the 5’ motif (site 1) and middle motif (site 2), indicating that ZNF410 binding to the 3’ motif facilitates its binding to the other motifs (Figures 5H and S6O). Furthermore, altering orientation of the 3 motifs (MT8) diminished all the ZNF410 binding, but inversing the whole peak5 had no effect (Figures 5I and S6N), in line with the results from luciferase assays (Figure 5G). More importantly, reversing the orientation of 3’-motif (MT11) also led to significant reduction of ZNF410 binding to the other sites, exhibiting a cooperative binding (Figures 5I and S6O). Taken together, these findings reveal functional collaborations among the ZNF410 sub-HCTs *in vivo*.

### The ZNF410 sub-HCTs within the peak5 controls the accessibility and activity of the whole distal enhancer

When mutating all three motifs in peak5, the ZNF410 binding across the whole distal enhancer harboring 11 ZNF410 motifs was markedly interrupted (Figure 5B), consequently causing reduced accessibility and enhancer activity (Figures 5D and 6A). Interestingly, when perturbing the distal enhancer with CRISPRi, we noted that sgRNAs targeting peak3 mainly diminishes the ZNF410 binding to peak3 and peak4 while sgRNAs targeting peak5 strikingly influenced ZNF410 binding to the whole distal region including peak3, peak4 and peak5 (Figures S2P and S2R). These observations indicate that ZNF410 binding to the distal enhancer appears to be regulated by the peak5. To confirm this idea, we performed luciferase assays with vectors containing individual or combination of these ZNF410 peaks. Notably, combining peak3, peak4 with peak5 exhibited synergistic effect on luciferase expression (Figure 6B). Importantly, incorporation of mutant peak5 lost ZNF410 binding led to 64%-83% reduction of luciferase activity while incorporation of mutant peak3 or peak4 only caused 23%-31% reduction (Figures 6C and 6D), compared with WT peaks, indicating that the ZNF410 binding to peak5 plays a major role on driving the activity of the distal enhancer, in line with the above data *in vivo*. To further reveal the mechanism, we generated two independent homozygous single-cell clones with deletion of peak5 (Figures S6P and S6Q), and then measured the ZNF410 binding, accessibility and H3K27ac marks of the *CHD4* enhancers (Figure 6A). Surprisingly, deletion of the peak5 had minimal effect on ZNF410 binding and chromatin accessibility of peak3 and peak4 (Figure 6A), distinct from that upon mutating the ZNF410 motifs of peak5. Peak5 deletion also displayed weaker effect on the H3K27ac levels at the region of peak3 and peak4 than that of peak5 motif mutation clones (Figure 6A). Lastly, compared with the combination of peak3 and peak4, addition of mutant peak5 severely impaired its luciferase activity (Figure 6E). These data together indicate that the ZNF410 sub-HCTs in peak5 act as “switch” to control the ZNF410 binding, and thus the accessibility, activity of the *CHD4* distal enhancer.

**Figure 6.**
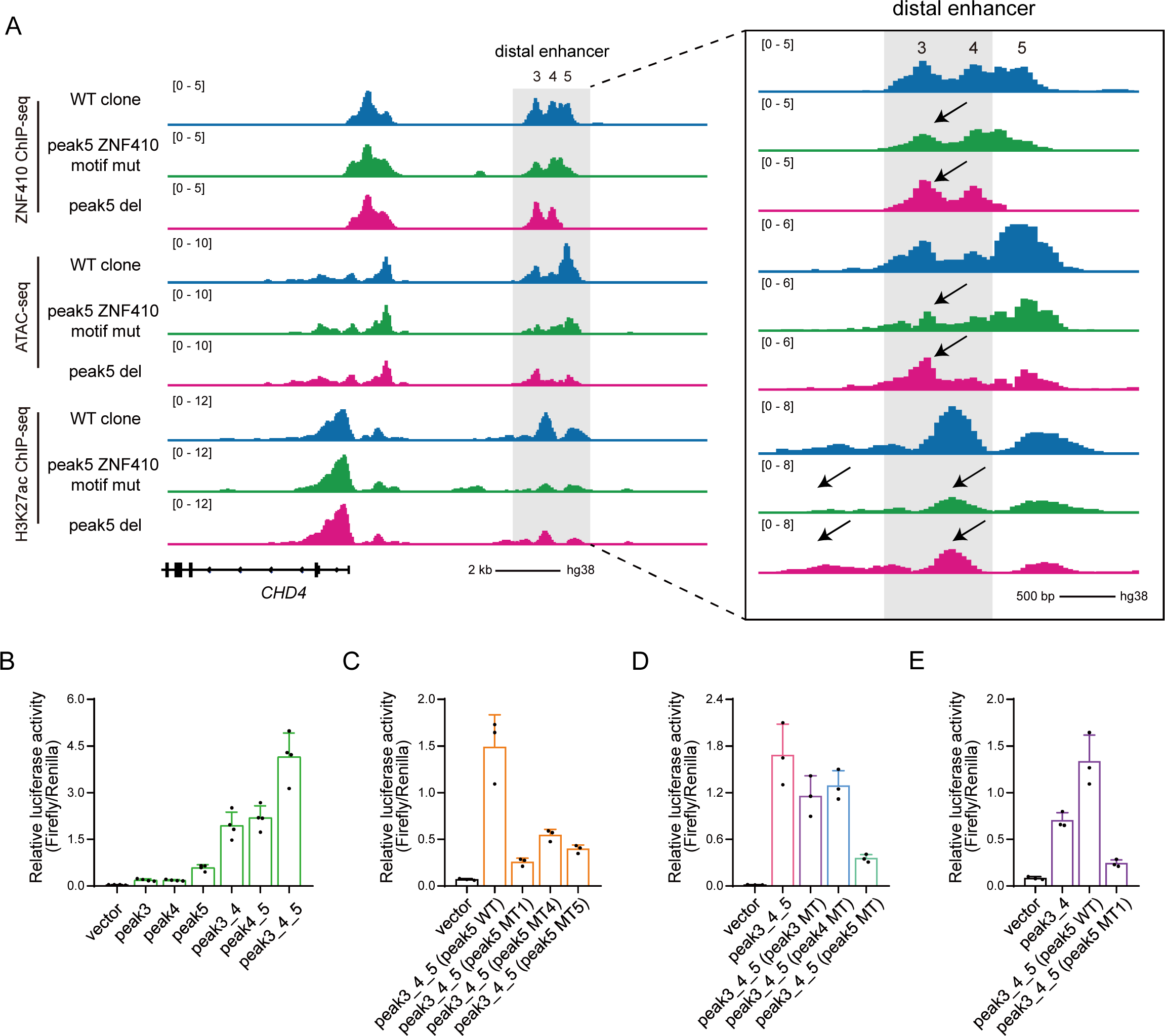
The function of peak5 sub-HCTs within the distal enhancer. (A) ChIP-seq profiles of ZNF410 and H3K27ac, as well as ATAC-seq tracks, in the *CHD4* distal enhancer regions of the CRISPR clones. (B) Results of luciferase reporter assays to measure the activity of ZNF410 peaks of the *CHD4* distal enhancer. Results are shown as mean ± SD (n=4). (C-E) Luciferase assays to measure the activity of the *CHD4* distal enhancer variants. Results are shown as mean ± SD (n=3), related to Figure S6.

## DISCUSSION

In this study, we attempt to dissect mechanisms of ZNF410 regulating *CHD4* gene expression given its uniqueness as a metazoan TF. Through profiling histone marks and chromatin accessibility we define that the *CHD4* regulatory regions are composed of two active enhancers. ZNF410 activates *CHD4* gene transcription via maintaining the chromatin accessibility of these *CHD4* enhancers. The two enhancers function to activate *CHD4* levels in an additive manner. Moreover, perturbation of the *CHD4* enhancers robustly elevates the γ-globin levels, a known target of CHD4/NuRD complex, in erythroid cells. Lastly, we identify the key nucleotides of ZNF410 motif for binding to DNA, and demonstrate that ZNF410 cooperatively binds to its sub-HCTs within the peak5 (Figure S6O), which act as “switch motifs” to control ZNF410 binding, the chromatin accessibility and activity of the whole distal enhancer (Figure S6R). These findings thus illustrate a novel mechanism through which TF cooperates to bind to DNA via homotypic clustered motifs for modulating gene transcription in cellular contexts, exposing a complicated functional hierarchy of motifs.

The bacterial CRISPR/Cas system has inspired the development of novel approaches to explore the functions of regulatory elements.^52^ To identify functional enhancers and relationships between enhancers, we deleted individually and combinatorially the *CHD4* enhancer regions by CRISPR DNA fragment editing and revealed that the two enhancer elements additively modulate *CHD4* gene expression. A previous study reveals that the *CHD4* expression decreases by 56%–79% after deletion of the 6.7 kb or 6.9 kb upstream elements in HUDEP-2 cells.^24^ Compared with those data, we only focused on the ZNF410-binding regions (approximately 2.5 kb sequences), with a less damage to DNA than in the other study, further indicating that the DNA elements between the proximal enhancer and distal enhancer have minimal effect on *CHD4* transcription (Figures 2C-2E). To avoid DNA cleavage by CRISPR/Cas9, CRISPRi was used to block the ZNF410 binding sites and thus inactivate the *CHD4* enhancer regions. Notably, the activity of *CHD4* enhancer regions were significantly impaired via dCas9-KRAB system. Nevertheless, both approaches consistently confirmed that ZNF410 binds to the two enhancers to additively activate *CHD4* transcription. What accounts for the exquisite specificity of ZNF410-*CHD4* regulation? When scanning the human genome dividing into 3-kb window with ZNF410 motif, only three windows comprised more than two ZNF410 motifs, including the *CHD4* proximal enhancer, distal enhancer, as well as the intron of *GALNT18*, and the vast majority of windows harbors one motif instance.^24^ Notably, only 6 sites with a single motif were bound by ZNF410, but no transcriptional output (Figures S1F and S1L). Moreover, mutating any two of the three ZNF410 motifs within the peak5 can completely abolish the luciferase expression (Figure 5A), consistent with *in vivo* results that a single motif is insufficient to activate transcription. Furthermore, despite presence of the ZNF410 HCTs within the intron of *GALNT18* locus, we demonstrate that variant A10C existed in each of the ZNF410 motifs within this HCTs eliminates luciferase activity of the peak5 (Figure S5D), presumably through disrupting ZNF410 binding to DNA, which could explain the absence of ZNF410 occupancy at this site. Lastly, both ZNF410 protein and its HCTs within the *CHD4* enhancer regions are highly conserved in mammals, suggesting a crucial role of this regulatory network. Therefore, based on these data, we propose that the cooperative binding of ZNF410 to its HCTs is required to drive transcription, which determines the unique regulation of ZNF410-*CHD4* axis during evolution.

Combinatorial TFs recognize specific DNA sequences to maintain chromatin accessibility and activate gene transcription.^1,53^ HCTs, a group of adjacent binding sites for the same TF, are present in both promoters and enhancer elements to influence gene expression.^20^ Several studies have reported that TF binds to its HCTs in an additive fashion. For instance, about 24 NF-κB binding clusters comprising three or more conserved NF-κB sites upstream of 23 genes are identified in the human genome. The transcription of NF-κB target genes gradually increases with increasing NF-κB nuclear concentrations. Remarkably, NF-κB binds independently to adjacent sites to promote additive RNA Pol II recruitment, indicating an additive but not cooperative manner among NF-κB binding sites in the clusters to initiate gene transcription.^54^ A separate study reveals that the transcriptional repressor ZEB1 binds to its HCTs in an additive manner as increasing ZEB1 concentrations results in an analog increase in occupancy of all ZEB1 binding sites *in vitro*.^55^ Only few studies have reported that the same TF collaborates to bind to DNA in bacterial or in *Drosophila*. The λ phage repressor binds cooperatively to the three sites in the right operator (O_R_).^56^ However, the binding pattern is alternate pairwise between repressor dimers, because when the O_R_ region is wild type, O_R_2 cannot interact simultaneously with dimers bound at O_R_1 and O_R_3.^56,57^ The *Drosophila* morphogenetic protein Bicoid is required for the formation of the *Drosophila* embryo along the anterior–posterior (A–P) axis. Bicoid binds to DNA with pairwise cooperativity and Bicoid bound to a strong site helps Bicoid bind to a weak site.^58,59^ These studies focus on the binding affinity of TF and verify *in vitro* by evaluating the relative concentration of repressor dimers^56^ or reporter gene expression.^58^ Recently, massively parallel reporter assays (MPRAs) has been utilized to study the function of HCTs,^60–62^ however, this approach may not recapitulate the truth at the native locus. We determined the cooperative binding of ZNF410 to the *CHD4* enhancer regions through both reporter assays and *in vivo* mutant motif knock-in clones that only replace 2 nucleotides within the ZNF410 motif to disrupt its binding.

Pioneer factors directly bind to the *cis*-regulatory elements to overcome the nucleosomal barrier and recruit multiple other TFs to facilitate nucleosome eviction.^50,63^ A recent study has shown that pioneer factor OCT4 binds to the nucleosome and facilitates cooperative binding of additional OCT4 and of SOX2 to their internal binding sites in a nucleosome-dependent manner via biochemistry and structural biology.^64^ Compared with OCT4, TF ZNF410 appears to be uncapable to bind to nucleosomal DNA *in vitro*. Meanwhile, we found that the 3’motif mutation has no obvious effect on the ZNF410 binding to additional sites in the naked DNA by EMSA assays. However, several evidences have suggested that a cooperative DNA-binding mechanism cannot be ruled out simply because TF does not bind cooperatively to naked DNA *in vitro*.^45,47,65,66^ The sub-HCTs of the ZNF410 peaks within the distal enhancer occupy less than 147 bp, likely locating in the same nucleosome. Cooperative ZNF410 binding to the sub-HCTs is likely involved in nucleosome eviction, further maintaining chromatin accessibility of the *CHD4* enhancers. Indeed, we showed that disabling the ZNF410 binding to the 3’ motif significantly impairs its binding to the other motifs of the peak5 sub-HCTs through episomal ChIP-qPCR in cells and ZNF410 ChIP-seq and ATAC-seq in mutant motif knock-in clones. Moreover, the motif at the 3’ ends of each peak of the distal enhancer controls the luciferase activity (Figures 5A and S6A). One possible explanation is that cofactors are required for ZNF410-mediated cooperative DNA binding.^65–70^ Further studies are needed to better understand what cofactors and how they act with ZNF410 to bind DNA to further regulate the chromatin accessibility and activity of the *CHD4* enhancers. Recent evidences have suggested that single genetic variants of TF binding sites can modulate the activities of multiple regulatory elements at kilobase and megabase resolution within topologically associated domains (TADs) and thus influence chromatin state and gene expression.^71–75^ A plausible explanation of these is that a single genetic variant might directly regulate the accessibility of a master enhancer region,^73,74^ exhibiting cooperation between enhancers. In contrast, we focus on the cooperation between motifs within TF ZNF410 bound enhancer. Importantly, the sub-HCTs within the peak5 of the distal enhancer exhibits “switch-like” effect on regulating the ZNF410 binding and chromatin accessibility of the whole enhancer region, thus resulting in an all-or-none occupancy of the HCTs within this enhancer. Therefore, we propose a model that ZNF410 initially binds to the sub-HCTs within the peak5 and then gains access to peak3 and peak4 (Figure S6R). Interestingly, unlike mutating the sub-HCTs of peak5, deleting this region has minimal effect on ZNF410 binding to peak3 and peak4. One plausible explanation is that deletion of the peak5 creates new genomic sequence without ZNF410 motifs so that the chromatin could still open from the 3’ end of peak4 to maintain the function of the distal enhancer (Figure S6R). Compared with the peak5 at 3’ end of the distal enhancer, the peak2 at 3’ end of the proximal enhancer is little conserved during mammalian evolution. Moreover, unlike peak5, perturbation of the ZNF410 binding in peak2 exhibits no effect on peak1 according to the ChIP-seq results (Figures S2O and S2R). Therefore, we speculate that there is likely no existence of such “switch motifs” in peak2 of the proximal enhancer. In sum, our findings uncover a paradigm regarding the usage of HCTs, exposing complicated functional hierarchy of motifs for regulating gene expression.

## ACKNOWLEDGMENTS

We thank all members of our laboratory for helpful comments and discussions. We also thank Yanhui Xu laboratory at the Fudan University for assistance with nucleosome reconstitution. We acknowledge the Medical Science Data Center in Shanghai Medical College of Fudan for providing data analysis platform. This work was supported by the National Natural Science

Foundation of China (32370614); Lingang Laboratory, China grant LG-QS-202204-05.

## AUTHOR CONTRIBUTIONS

S.X., G.A.B., and X.L. conceived the study and designed the experiments. S.X. carried out all the cell-based experiments. S.X., R.R., and X.C. performed DNA binding assays. L.C.L. and M.C. carried out the band retardation experiments. H.L. and B.X. helped with the experiments. S.X., and C.P. carried out luciferase assays. S.X. analyzed RNA-seq, ChIP-seq, and ATAC-seq data. S.X. and X.L. wrote the manuscript with input from all authors.

## Data and code availability

The RNA-seq, ChIP-seq, and ATAC-seq data are accessible through GEO series accession number GSE256315, GSE256316, and GSE256317, respectively.

## Methods and Materials

### Cell culture

K562 cells were cultured in IMDM supplemented with 10% fetal bovine serum and 1% penicillin-streptomycin solution. HUDEP-2 cells were expanded and differentiated as described.^76^ Briefly, cells were expanded in StemSpan SFEM supplemented with 1 µg/ml doxycycline, 1 µM dexamethasone, 50 ng/ml SCF, 3 IU/ml erythropoietin and 1% penicillin-streptomycin solution at a maximum density of 1 × 10^6^ cells/ml. HUDEP-2 cells were differentiated with IMDM media with 320 µg/ml transferrin, 5% fetal bovine serum, 2% penicillin-streptomycin, 50 ng/ml SCF, 3 IU/ml erythropoietin, 10 µg/ml insulin, 10 μg/ml heparin and 1 µg/ml doxycycline for 7 days. HEK293T cells, U87 cells, U251 cells, and COS-7 cells were maintained in DMEM media with 10% fetal bovine serum and 1% penicillin-streptomycin solution. BT-474 cells were maintained in RPMI 1640 media with 10% fetal bovine serum and 1% penicillin-streptomycin solution. SK-BR-3 were cultured in McCoýs 5A media with 10% fetal bovine serum and 1% penicillin-streptomycin solution. All cells were cultured at 37°C with 5% (v/v) of CO_2_.

### Plasmid construction

To prepare sgRNA plasmids for CRISPR DNA fragment editing, complementary oligonucleotide pairs were annealed to generate dsDNA with 5’ overhangs of CACC and AAAC. The annealed dsDNA was sub-cloned into the BsmbI-digested lentiviral sgRNA-EFS-GFP/mCherry vector under the control of U6 promoter (LRG, Addgene #65656). To improve the transcription efficiency, an additional 5’ G nucleotide was added to all the sgRNA oligonucleotide designs that did not begin with a 5’ G.

To knock in the AA to GG of the ZNF410 motifs, the left and right homologous arm were amplified by PCR from genomic DNA for CRISPR mediated homologous recombination. Two homologous arms, the ZNF410 motif mutant sequences, and the linearized pcDNA3 vector, digested by NheI and BamHI were jointed together with 15-20 bp overlapping sequences using the NEBuilder HiFi DNA Assembly Cloning Kit (NEB).

For luciferase assays, each WT ZNF410 peak was amplified by PCR from genomic DNA and cloned into pGL4.24 vector (Promega, #E8421), digested by KpnI and XhoI. To explore the influence of cooperation between ZNF410 motifs and the orientation of ZNF410 motifs, all the variants of ZNF410 motif mutant sequences were synthesized and ligated into pGL4.24 vector, digested by KpnI and XhoI.

For electrophoretic mobility shift assay (EMSA), the ZNF410 full length and ZNF410-ZF variant were sub-cloned into pcDNA3 vector. For protein expression and purification, the ZNF410 full length and ZNF410-ZF variant were sub-cloned into pGEX-6P-1 vector. The plasmids constructed were confirmed by Sanger sequencing and primers are listed in Supplemental Table S2.

### Lentiviral transduction

Lentivirus was produced by transfecting HEK293T cells for each stable cell line construction. Briefly, about 10 μg lentivirus transfer plasmid, 7.5 μg packaging helper plasmid (psPAX2, Addgene #12260), 5 μg envelope helper plasmid (pMD2.G, Addgene #12259) and 80 μl of 1mg/ml polyethylenimine (PEI) were mixed and incubated for 15 mins. The mixture was added to HEK293T cells grown in 10 cm plates to 80-90% confluence. After 8 hr, the medium was replaced with fresh complete medium. The virus was collected after transfection for 24 hr and 48 hr. For K562 and HUDEP-2 cells infection, the mixture of virus, medium, 8 μg/ml polybrene and 10 mM HEPES was spun at 2250 rpm for 1.5 hr at room temperature. Infected K562 and HUDEP-2 cells were selected for mCherry+, or GFP+, or mCherry+ and GFP+ cell sorting after 48 hr.

### Knock out and knock in by CRISPR DNA fragment editing

K562 cells were transfected with the CRISPR gene editing components, consisting of 1.5 μg Cas9 plasmid, 1.5 μg dual sgRNA plasmids or 1 μg sgRNA plasmids and 1 μg homologous recombination donor using Neon™ NxT Electroporation System (Invitrogen). After 48 hr, the culture medium containing 1 μg/ml of puromycin was used for 4 days of drug selection. After identifying the mixture genotyping by PCR, the cells were diluted and plated into 96-well culture plates. All the single-cell cells were cultured about 10 days, and then were screened for target clones by PCR. Two or three independent single-cell homozygous clones were obtained and the genotype of single-cell clones was confirmed by Sanger sequencing. Primers are listed in Supplemental Table S3.

### RNA extraction and RT-qPCR

Total RNA was extracted using the FastPure Cell/Tissue Total RNA Isolation Kit V2 (Vazyme). According to the manufactureŕs instructions, about 500 ng total RNA was used in reverse transcription (Vazyme). In the process of the reverse transcription, gDNA wiper was used to remove genomic DNA. The cDNA template was diluted and used for qPCR with SYBR Green master mix (Roche). After normalization with *GAPDH*, the relative expression levels of each gene were calculated by 2^-ΔΔCt^ method. All qPCR experiments were performed with at least two biological replicates. Data statistical tests were performed with GraphPad Prism (v8.0). Primers for qPCR are listed in Supplemental Table S4.

### Luciferase assays

Each ZNF410 peak and all the variants of ZNF410 motif mutant sequences were sub-cloned into pGL4.24 plasmid (Promega, #E8421). About 1.2 μg WT or constructed pGL4.24 plasmid and 0.2 μg pGL4.74 control plasmid (Promega, #E6921) were co-transfected into K562 cells by Neon™ NxT Electroporation System (Invitrogen). After 24 hr transfection, the relative luciferase activity was measured according to manufactureŕs instructions (Promega, #E1910). The relative luciferase activity was calculated by the ratio of firefly luciferase activity to internal Renilla luciferase activity.

### Protein expression and purification

The full-length of human ZNF410 (NP_001229855.1) and five zinc finger domains ZF1-5 (residues 217-366) of ZNF410 were cloned into pGEX-6P-1 vector with a GST fusion tag. The expression plasmids were transformed into Escherichia Coli strain BL21-Codonplus(DE3)-RIL (Stratagene). The expression and purification of each protein was performed as described previously.^77^ Briefly, bacteria were grown in LB broth in a shaker at 37°C until the optical density (OD_600_) reached ∼0.8, the protein expression was induced by adding 0.2 mM isopropyl-β-D-thiogalactopyranoside (IPTG) with subsequent growth for 20 hr at 16°C. Cells were collected, sonicated and GST-fusion proteins were purified with GSTrap column. The GST fusion were digested with PreScission protease to remove the GST fusion tag. The protein was frozen and stored at −80°C.

### Electrophoretic mobility shift assay (EMSA)

About 5 μg empty pcDNA3 vector or ZNF410 full-length and ZNF410-ZF plasmids was transiently transfected with FuGENE 6 (Promega) into 10 cm plates of COS-7 cells. Cells were harvested after 48 hr transfection and nuclear extracts were prepared as previously described.^78^ The two strands of short probe was labeled with [γ-^32^P]-adenosine triphosphate (PerkinElmer). After that, the annealed probes were purified with Quick Spin Columns for radio-labeled DNA Purification (Roche). ZNF410 full-length or ZNF410-ZF were overexpressed and harvested from COS-7 cells, and a ‘COS empty’ extract was used as negative control. The gels were exposed with a FUJIFILM BAS Cassette2 phosphor screen overnight and scanned by the Typhoon FLA 9500 Laser Scanner on the second day.

For the binding of purified ZNF410 ZF1-5 and FL with linear DNA, the DNA probe was amplified by PCR using 5′ biotin-labeled primers. EMSA was performed with the LightShift Chemiluminescent EMSA Kit according to the manufactureŕs protocols (Thermo Fisher Scientific). The biotin-labeled DNA probe was incubated with purified protein for 20 mins at room temperature. The binding reactions were electrophoresed on 6% native polyacrylamide gels in 0.5 × TBE buffer at 80 V for 100-120 mins at 4°C and transferred to a nylon membrane. After crosslinking with UV-light for 1 min, the membrane was blocked in the Blocking Buffer for 15 mins with gentle shaking and incubated with Stabilized Streptavidin-Horseradish Peroxidase Conjugate for additional 15 mins with gentle shaking. After that, the membrane was washed four times by Wash Buffer and incubated with Substrate Equilibration Buffer for 5 mins, followed by incubation in the Substrate Working Solution for 5 mins. The membrane was scanned by the ChemiDoc Imaging System (Bio-Rad).

### Nucleosome reconstitution

Nucleosome reconstitution was performed according to previously published protocols.^79^ The histone proteins (H2A, H2B, H3 and H4) were expressed in *E. coli* and extracted from inclusion bodies and refolded to form histone octamer through dialysis. The 200 bp DNA fragment was amplified by PCR. For the assembly, the histone octamer and purified DNA fragment were mixed in 1:1 ratio, and the nucleosome was reconstituted by dialyzed gradually about 24 hr. The 200 bp DNA fragment sequence for assembly is: 5’-CCCATAATACTCCCTTCCACCACCACCCCCCCTTCCCGCCCCGCACCCCTTCCTCCCG CCTCCAGTTGGAGCCGGGCCTCACCAACCCATAATATTCCCCAGTCTCCTAAGGCTGTTCCAGCCCATAATACTCTCCCTGTCTCCTAAGGCCTCTCCATCCCATAATACTGCCAGACGCCTGACCTGTGGGCCAAACCCCA-3’. The samples were on a 6% native polyacrylamide gel, followed by stained with SYBR Gold dye (Thermo Fisher Scientific).

### DNA-binding assay

Fluorescence polarization (FP) method was used to measure the binding affinity with Synergy 4 Microplate Reader (BioTek). Aliquots (5 nM) of 6-carboxy-fluorescein (FAM)-labeled ZNF410 motif and ZNF410 mutant motif was incubated with varied amount of proteins (0-2.5 mM) in 20 mM Tris-HCl, pH 7.5, 300 mM NaCl, 5% glycerol and 0.5 mM tris-(2-carboxyethyl)phosphine for 10 min at room temperature before measurement. The data were processed with equation [mP] = [maximum mP] × [C] / (K_D_ +[C]) + [baseline mP], in which mP is millipolarization and [C] is protein concentration. The K_D_ value for each protein–DNA interaction was derived from two replicated experiments. The data were processed using GraphPad Prism (v8.0).

### Immunoblot analysis

For the purpose of the total protein extraction, about 1-2 × 10^6^ cells were lysed with the RIPA lysis buffer (Thermo Fisher Scientific) with 1 × protease inhibitors (Roche) and 1 × phenylmethylsulfonyl fluoride (PMSF) for 30 mins on ice. Protein lysate was denatured at 95°C for 5-10 mins. Next, about 20-30 μg proteins were separated by SDS-PAGE and transferred to nitrocellulose membranes. The membranes were blocked by 5% nonfat milk in TBST and incubated with a primary antibody overnight at 4°C. The membranes were washed with 1 × TBST three times, followed by incubation at room temperature for 1 h with the secondary antibody, and scanned by the ChemiDoc Imaging System (Bio-Rad).

### RNA-seq

Total RNA was extracted as described above. RNA-seq experiments were performed using VAHTS Universal V6 RNA-seq Library Prep Kit for Illumina according to the manufacturés protocol (Vazyme). mRNA was enriched from 1 μg of total RNA using mRNA capture beads and fragmented by heating. Random primers were used in reverse transcription of the first-strand cDNA and synthesis of the second-strand cDNA reaction. After that, cDNA was selected and purified by AMPure XP beads (Beckman). The purified cDNA was end-repaired and then ligated with VAHTS RNA adapters. The ligated cDNA was amplified with the PCR program for 13 cycles. The size and quality of each library were evaluated by Bioanalyzer 2100 (Agilent Technologies). All RNA-seq libraries are sequenced on an Illumina NovaSeq 6000 platform. All RNA-seq experiments were performed with two biological replicates.

### Chromatin immunoprecipitation (ChIP)

About 1-2 × 10^7^ cells were cross-linked with 1% formaldehyde at room temperature for 10 mins, followed by quenching with glycine for 5 mins. Cells were resuspended in the ice-cold cell lysis buffer (10mM Tris pH 8, 10 mM NaCl, 0.2% Igepal) and nuclei lysis buffer (50 mM Tris pH 8, 1% SDS, 10 mM EDTA) with 1 × PMSF and 1 × protease inhibitors for 10 mins with slow rotations, respectively. The nuclei were sonicated for 30 mins at 4°C (a train of 30 s on and 30 s off for 30 cycles). After removal of the insoluble debris, the cell lysate was pre-cleared with 50 μl Protein A/G agarose beads for 2 hr at 4°C. Taking 4% of precleared chromatin set aside as input and the remainder was added Protein A/G beads pre-bound with specific antibodies against ZNF410 (Proteintech, Cat. # 14529-1-AP), H3K27ac (Abcam, Cat. # ab4729), H3K4me3 (Abcam, Cat. # ab8580) and H3K9me3 (Abcam, Cat. # ab8898) overnight. On the second day, the beads were washed once with ChIP wash buffer I (20 mM Tris pH 8, 50 mM NaCl, 2 mM EDTA, 1% Triton X-100, 0.1% SDS), twice with high salt buffer (20 mM Tris pH 8, 0.5 M NaCl, 2 mM EDTA, 1% Triton X-100, 0.01% SDS), once with wash buffer II (10 mM Tris pH 8, 0.25 M LiCl, 1 mM EDTA, 1% Igepal, 1% sodium deoxycholate) and twice with TE (10 mM Tris pH 8, 1 mM EDTA pH 8). Then, the complexes were eluted twice with 100 ul elution buffer. And then, NaCl and RNase A (Thermo Fisher Scientific) were added to samples as well as inputs and put at 37°C for 30 mins. After that, proteinase K (Thermo Fisher Scientific) was added and samples were incubated at 65°C overnight. The DNA was purified with phenol-chloroform, followed by ethanol precipitation, and then used for qPCR and DNA library preparation. All ChIP experiments were performed with two biological replicates.

### ChIP-qPCR and ChIP-seq

For ChIP-qPCR, the purified DNA was mixed with SYBR Green master mix (Roche) and specific primers. ChIP–qPCR primers are listed in Supplemental Table S5. For ChIP-seq, DNA library was constructed according to the instructions of VAHTS Universal DNA Library Prep Kit for Illumina (Vazyme). Briefly, the purified DNA was end-repaired and then ligated with adapters. After purified by AMPure XP beads (Beckman), the DNA was amplified with the PCR program. The size and quality of each library were evaluated by Bioanalyzer 2100 (Agilent Technologies). All ChIP-seq libraries were sequenced on an Illumina NovaSeq 6000 platform.

### Episomal ChIP-qPCR

Episomal ChIP-qPCR was performed as description with some modifications.^41^ Briefly, about 5 million K562 cells were transfected with 5 μg luciferase reporter plasmid using Neon™ NxT Electroporation System (Invitrogen). Cells were fixed after 24 hours and ChIP performed as described above. Primers for qPCR were designed to distinguish between episomal and endogenous DNA.

### ATAC-seq

ATAC-seq (assay for transposase-accessible chromatin using sequencing) was performed using the Omni-ATAC protocol with some modifications.^80^ Briefly, about 5 × 10^4^ K562 cells were lysed with resuspension buffer (10 mM Tris-HCl pH 7.4, 10 mM NaCl, 3 mM MgCl_2_) containing 0.1% NP-40, 0.1% Tween-20 and 0.01% digitonin. The lysate was then washed and incubated with transposition buffer (Vazyme) for 30 mins at 37 °C. After purified by AMPure XP beads (Beckman), the DNA was amplified with the PCR program. The size and quality of each library were evaluated by Bioanalyzer 2100 (Agilent Technologies). All ATAC-seq libraries were sequenced on an Illumina NovaSeq 6000 platform. All ATAC-seq experiments were performed with two biological replicates.

### RNA-seq data processing and analysis

For RNA-seq experiments, raw FASTQ files were aligned to the human reference genome (GRCh38/hg38) using HISAT2 (v2.2.1)^81^ with default parameters. SAMtools (v1.9)^82^ was applied to transform SAM into the BAM format. HT-seq (v0.6.1)^83^ was used to calculate the read counts of each transcript. RSEM (1.3.2)^84^ was used to calculate the expression levels of each transcript by FPKM (fragments per kilobase of exon per million fragments mapped). For differential expression analysis, we used adjusted *P* < 0.05 and log2 fold-change > 1 as thresholds to identify differentially expressed genes by DESeq2 package^85^ (1.36.0) in R (4.2.1). Commonly changed genes in both independent sgRNAs were considered to be significant. Data statistical tests were performed with GraphPad Prism (v8.0).

### ChIP-seq data processing and analysis

For ChIP-seq experiments, raw FASTQ files were aligned to the human (GRCh38/hg38) reference genome using Bowtie 2 (v2.2.5)^86^ with the default parameters. SAMtools (v1.9)^82^ was applied to transform SAM into the BAM format and removed PCR duplicates and unmapped reads. DeepTools (v3.5.1)^87^ was applied to transform BAM into the bigwig format, normalized using the counts per million (CPM), followed by visualizing in the Integrative Genomics Viewer (IGV).

The ZNF410 peak calling was done using MACS2 (v2.1.4)^88^ with the default parameters. The final peaks were those overlapped by both ZNF410 biological replicates, created by BEDTools (v2.30.0),^89^ was used for *de novo* motif analysis with the HOMER tool (v4.11).^90^ Read coverage was determined with BEDTools^89^ and normalized using CPM.

### ATAC-seq data processing and analysis

For ATAC-seq experiments, raw FASTQ files were aligned to the human (GRCh38/hg38) reference genome using Bowtie 2 (v2.2.5)^86^ with the default parameters. SAMtools (v1.9)^82^ was applied to transform SAM into the BAM format and removed PCR duplicates and unmapped reads. MACS2 (v2.1.4)^88^ was used to call ATAC-seq peak with the default parameters. Signal tracks were normalized using the CPM options in deepTools (v3.5.1)^87^ in order to create coverage bigwigs, followed by visualizing in the IGV. Read coverage was determined with BEDTools^89^ and normalized using CPM.

### Other statistical analysis

Results were presented as the mean ± SD. All statistical analyses were performed with GraphPad Prism (v8.0). Statistical significance was determined using unpaired Student’s *t*-test. *P* ≤ 0.05 was represented as ‘*’, *P* ≤ 0.01 was represented as ‘**’ and *P* ≤ 0.001 was represented as ‘***’, respectively.

---

**Figure S1.**
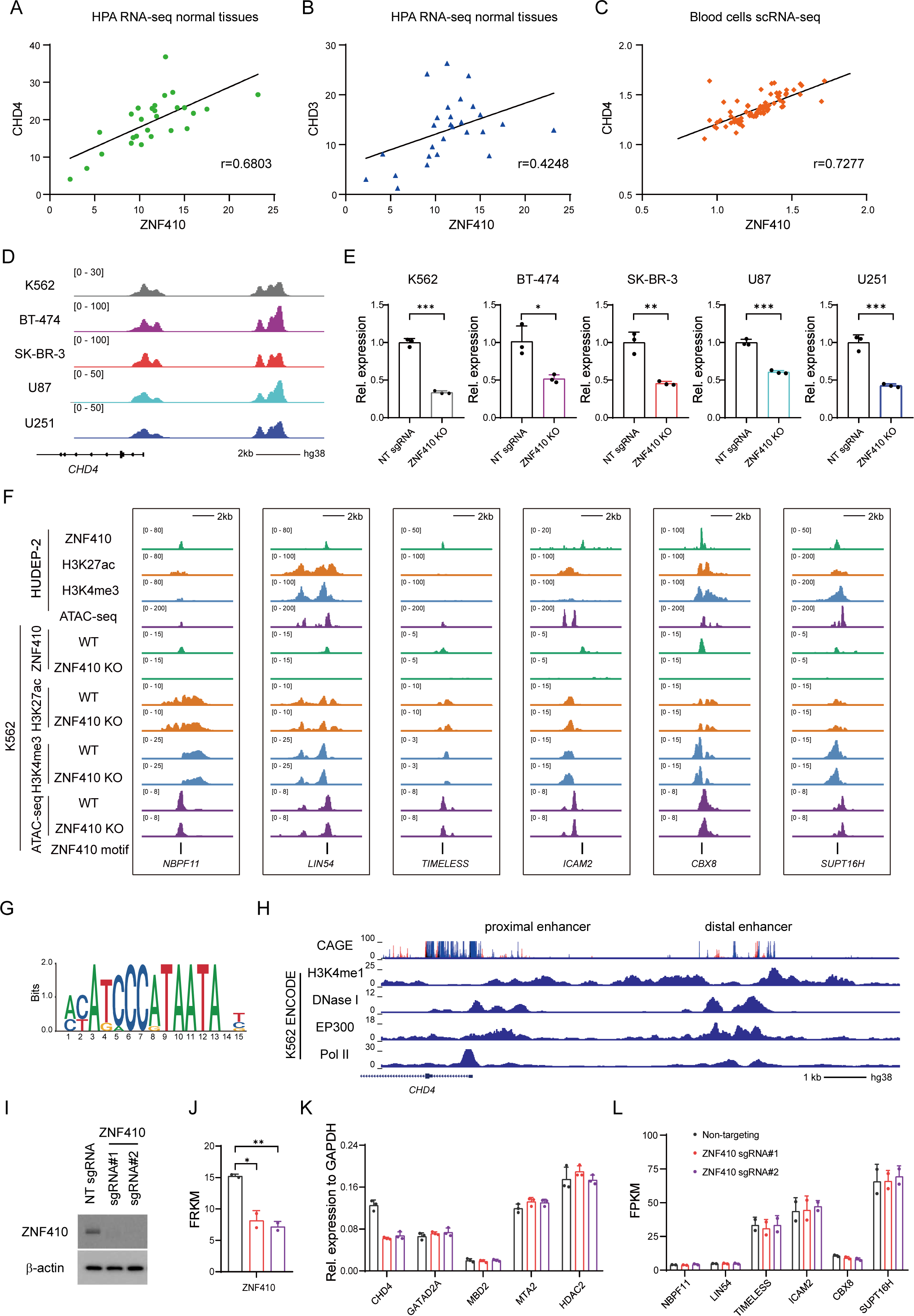
The pattern of ZNF410 regulating *CHD4* expression is highly similar between K562 and HUDEP-2 cells, related to Figure 1. (A) Correlation of *ZNF410* and *CHD4* mRNA levels across 27 human normal tissues. Data were obtained from the Human Protein Atlas (HPA) database. Expression level was calculated by RPKM (Reads per kilo base per million mapped reads). (B) Correlation of *ZNF410* and *CHD3* mRNA levels across 27 human normal tissues. (C) ChIP-seq profiles of the *CHD4* locus in multiple cell lines. (D) RT-qPCR analysis of *CHD4* in multiple ZNF410-depleted cell lines. Results are shown as mean ± SD (n=3). \**P* < 0.05, \*\**P* < 0.01, \*\*\**P* < 0.01; unpaired Student’s *t*-test. NT, non-targeting; KO, knockout. (E) Correlation of *ZNF410* and *CHD4* mRNA levels across blood cells. Single-cell RNA-seq (scRNA-seq) data were obtained from the Tabula Sapiens database. (F) ChIP-seq and ATAC-seq profiles of the *NBPF11*, *LIN54*, *TIMELESS*, *ICAM2*, *CBX8*, *SUPT16H* loci in HUDEP-2 and K562 cells. Upper panel: shown are ZNF410 (green), H3K27ac (orange), H3K4me3 (light blue) ChIP-seq, and ATAC-seq (purple) in HUDEP-2 cells. Lower panel: anti-ZNF410, anti-H3K27ac, anti-H3K4me3 (light blue) ChIP-seq, and ATAC-seq tracks of wild-type (WT) and ZNF410-depleted K562 cells. (G) The *in vivo* (ChIP–seq) binding motifs for ZNF410 in K562 cells by HOMER. (H) ChIP-seq profiles of the *CHD4* enhancer regions from the cap analysis of gene expression (CAGE) and ENCODE databases. (I) Immunoblot analysis using whole-cell lysates from K562 cell pools transduced with the indicated sgRNAs. (J) Expression levels of ZNF410 by RNA-seq analysis of WT and ZNF410-depleted K562 cells. Results are shown as mean ± SD (n=2). \**P* < 0.05, \*\**P* < 0.01; unpaired Student’s *t*-test. FPKM, Fragments per kilo base per million mapped reads. (K) RT-qPCR analysis of *CHD4*, *GATAD2A*, *MBD2*, *MTA2*, *HDAC2* in WT and ZNF410-depleted K562 cells. Results are shown as mean ± SD (n=3). (L) Expression levels of *NBPF11*, *LIN54*, *TIMELESS*, *ICAM2*, *CBX8*, *SUPT16H* by RNA-seq analysis of WT and ZNF410-depleted K562 cells. Results are shown as mean ± SD (n=2).

**Figure S2.**
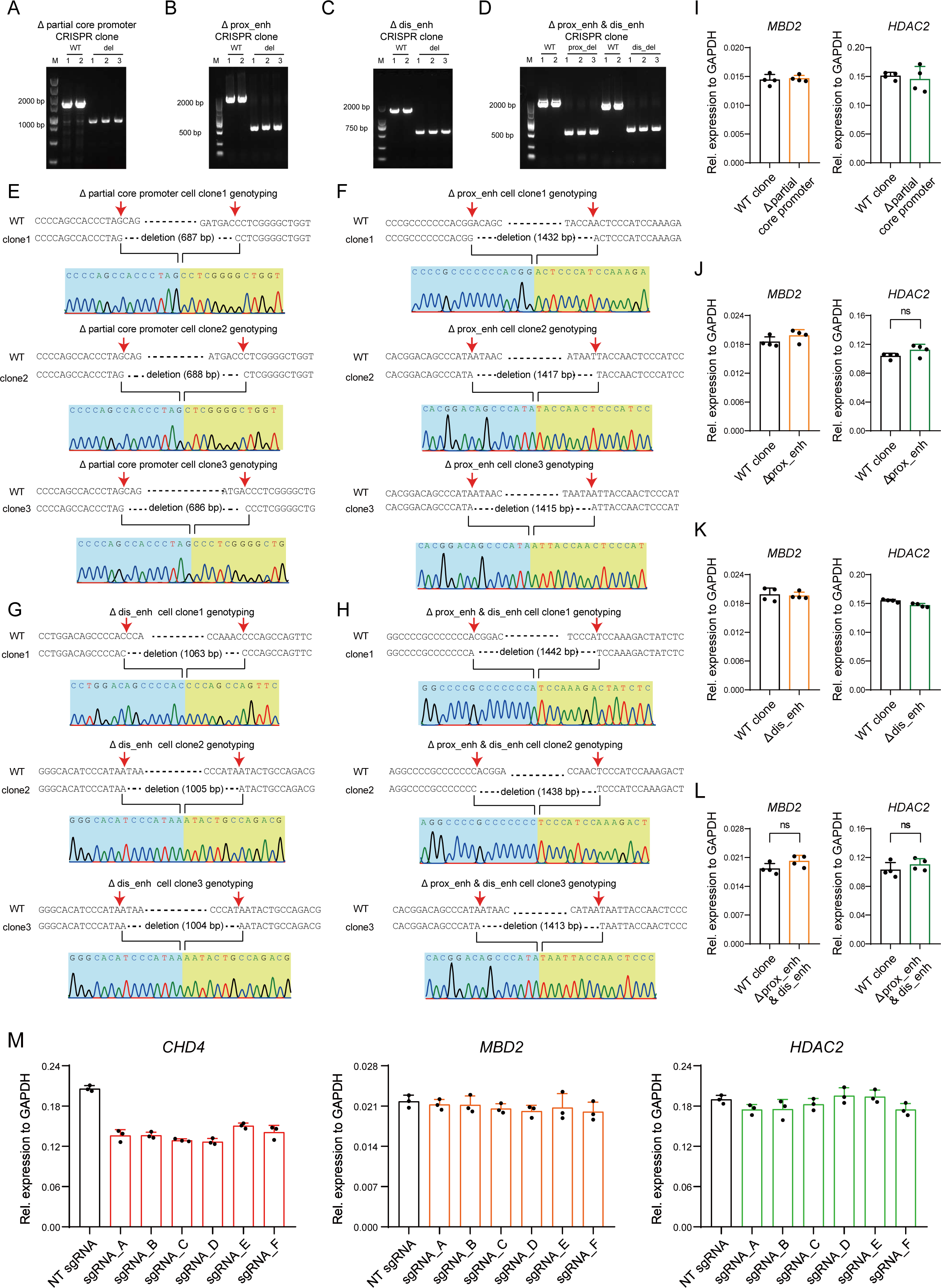
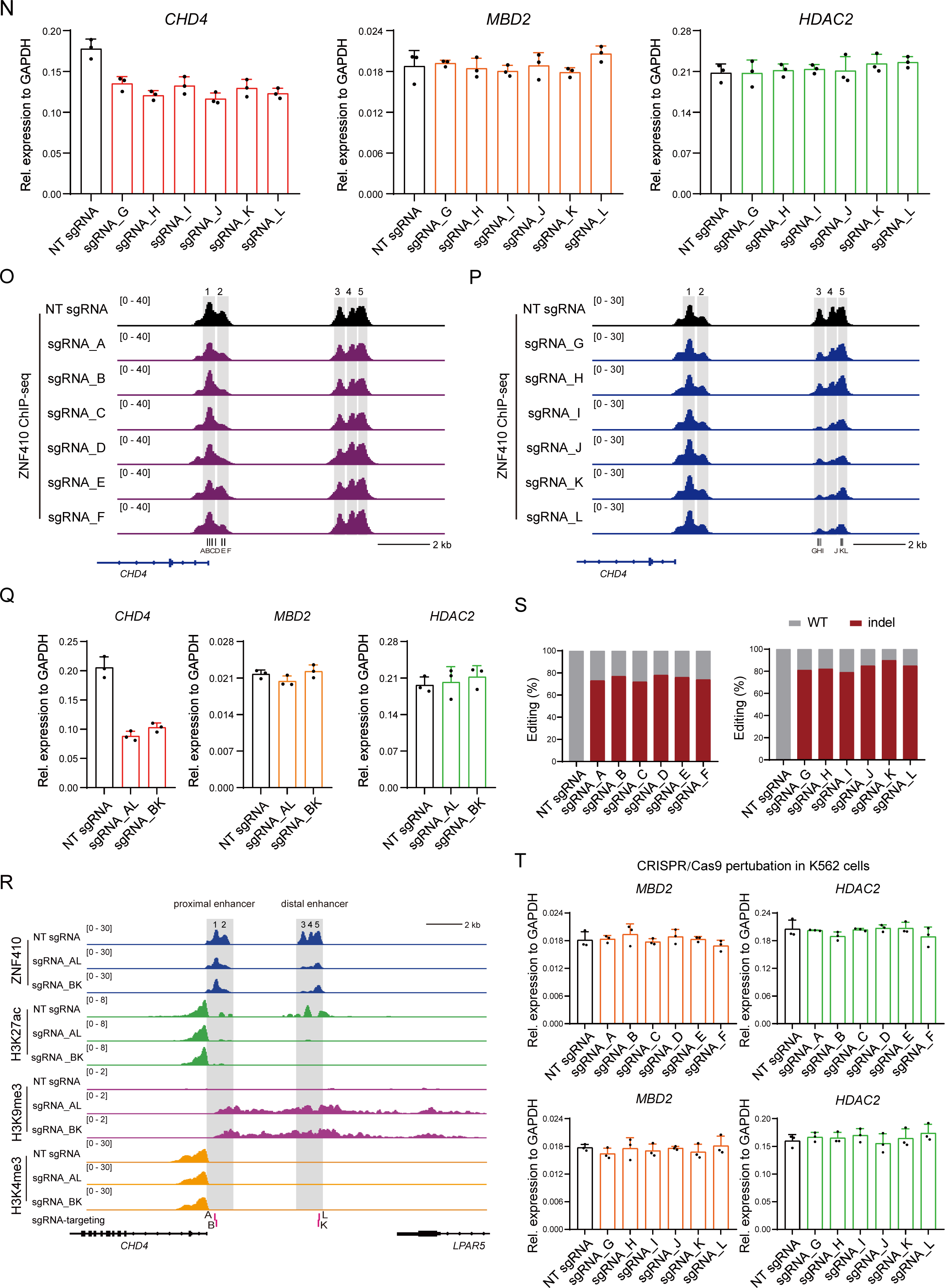
CRISPR genetic and epigenetic perturbation of ZNF410 motifs in the *CHD4* **proximal and distal enhancers in K562 cells, related to** Figure 2. (A-D) Genotyping of the homozygous single- and double-enhancer knockouts single-cell CRISPR clones. Shown are PCR products of partial core promoter deletion clones (A), single proximal enhancer (prox_enh) deletion clones (B), single distal enhancer (dis_enh) deletion clones (C) and double-enhancer (prox_enh & dis_enh) deletion clones (D), respectively. (E-H) Sanger sequencing traces of the homozygous single- and double-enhancer knockouts single-cell CRISPR clones. (I-L) RT-qPCR analysis of *MBD2* (orange) and *HDAC2* (green) in WT clone (n=4) and single- and double-enhancer knockouts single-cell clones (n=4). Results are shown as mean ± SD. ns, not significant. (M-N) RT-qPCR analysis of *CHD4* (red), *MBD2* (orange), and *HDAC2* (green) mRNA levels of K562 dCas9-KRAB stable cell lines with the distinct sgRNAs targeting the proximal enhancer and/or distal enhancer. Results are shown as mean ± SD (n=3). NT, non-targeting. (O-P) ChIP-Seq profiles of ZNF410 occupation in the *CHD4* proximal enhancer (O) or distal enhancer (P). (Q) RT-qPCR analysis of *CHD4* (red), *MBD2* (orange), and *HDAC2* (green) using pairs of sgRNAs simultaneously targeting the *CHD4* proximal and distal enhancers upon CRISPRi perturbation. (R) ChIP-seq profiles of ZNF410 (dark blue), H3K27ac (green), H3K9me3 (purple), and H3K4me3 (orange) in the *CHD4* proximal and distal enhancers upon CRISPRi perturbation. (S) Verifying the on-targeting sgRNA editing efficiency. Genomic DNA were extracted and subject to PCR and Sanger sequencing analysis. (T) *MBD2* (orange) and *HDAC2* (green) levels measured by RT-qPCR in each ZNF410 binding motif disruption of K562 cell pools via CRISPR/Cas9. Results are shown as mean ± SD (n=3). CRISPR/Cas9 pertubation in HUDEP-2 cells

**Figure S3.**
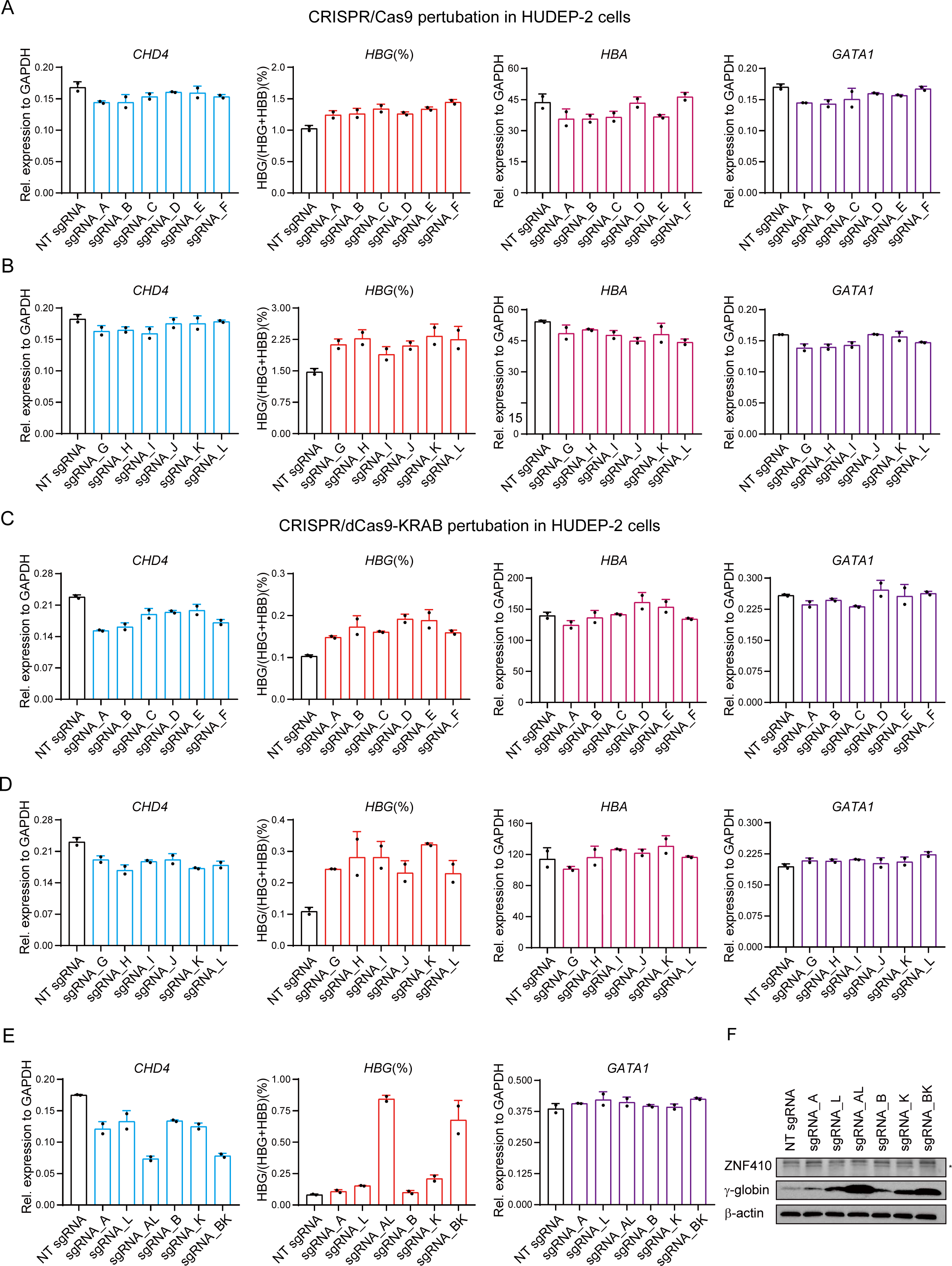
Perturbation of the *CHD4* enhancers robustly elevates γ-globin levels in HUDEP-2 cells. (A-B) Expression levels of *CHD4* (light blue), the ratio of *HBG*/(*HBG* + *HBB*) (red), α-globin (dark red), and *GATA1* (purple) by CRISPR DNA fragment editing. Results are shown as mean ± SD (n=2). NT, non-targeting. (C-D) Expression levels of *CHD4* (light blue), the ratio of *HBG*/(*HBG* + *HBB*) (red), α-globin (dark red), and *GATA1* (purple) upon CRISPRi perturbation. Results are shown as mean ± SD (n=2). (E) RT-qPCR analysis of *CHD4* (light blue), and the ratio of *HBG*/(*HBG* + *HBB*) (red), and *GATA1* (purple) mRNA levels using pairs of sgRNAs simultaneously targeting the *CHD4* proximal and distal enhancers. Results are shown as mean ± SD (n=2). (F) Immunoblot analysis using whole-cell lysates from differentiated HUDEP-2 cell pools transduced with the indicated sgRNAs. *, non-specific band.

**Figure S4.**
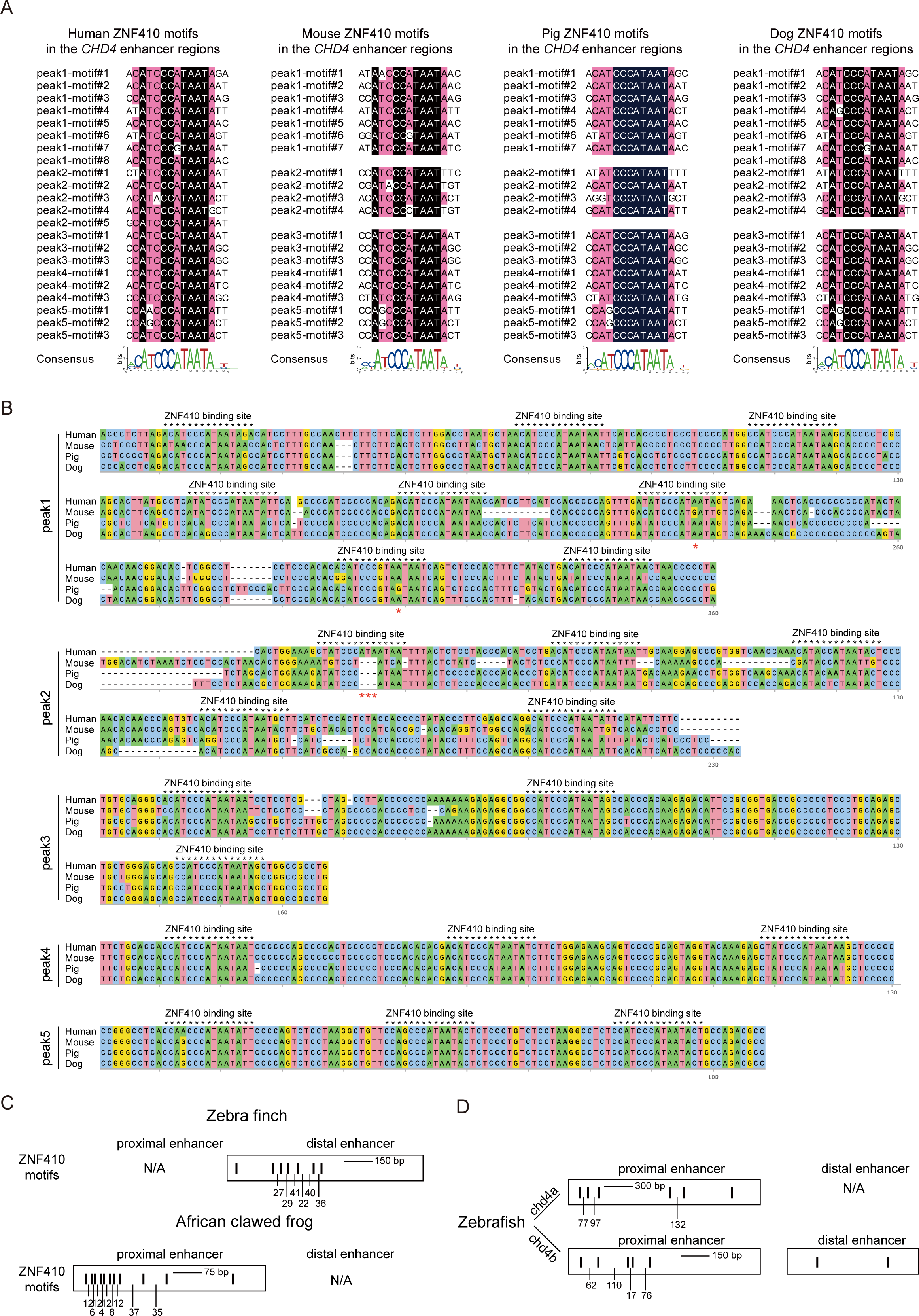
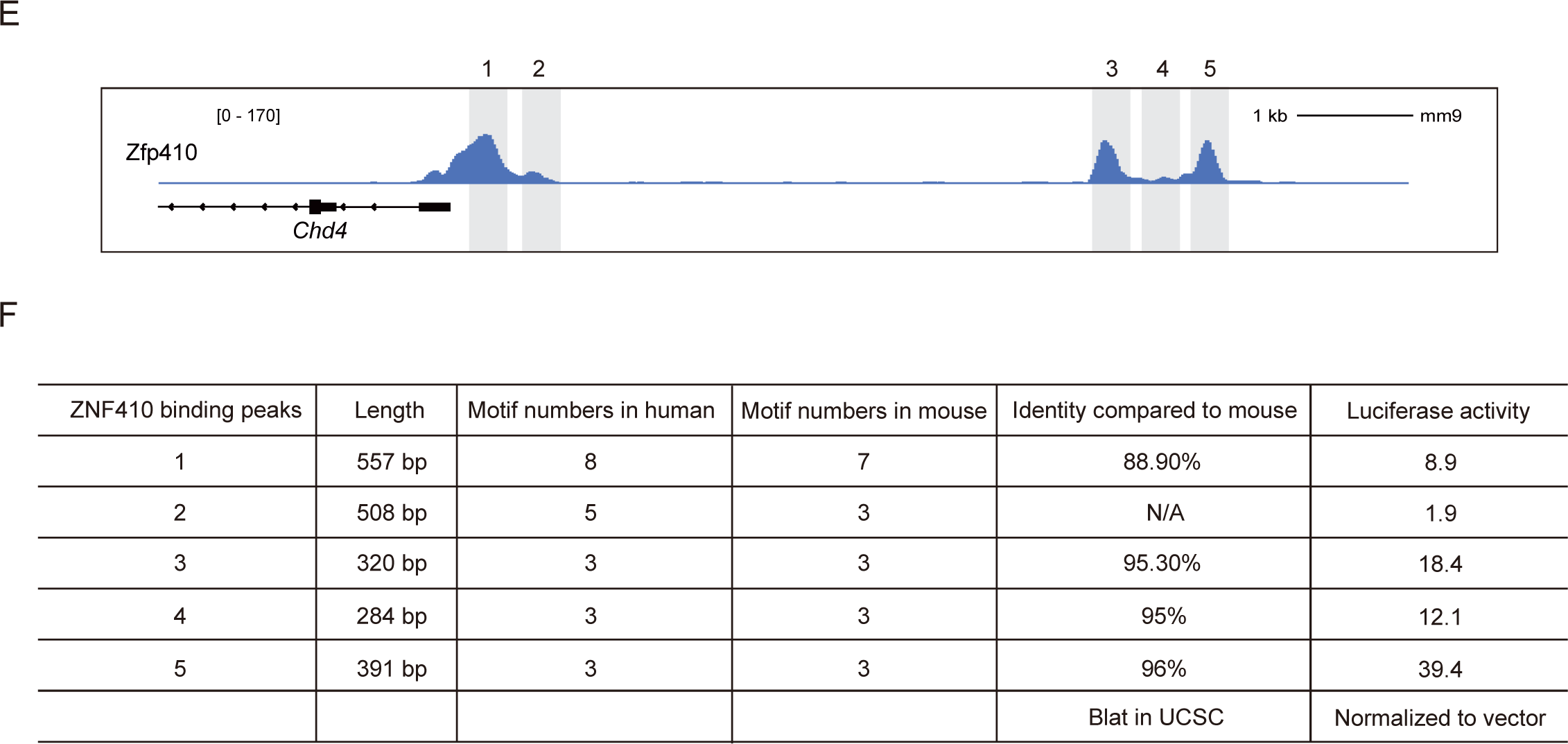
ZNF410 clustered motifs are conserved in mammals, related to Figure 3. (A) ZNF410 orthologs binding motifs in the *CHD4* enhancers. Black square corresponds to 100% identity and pink square means to 75% identity. (B) Multiple sequence alignment of ZNF410 binding regions of each ZNF410 peak among human, mouse, pig and dog. Position of the ZNF410 binding sites were highlighted with black asterisk. Mutated ZNF410 binding motifs were highlighted with red asterisk. (C) ZNF410 orthologs binding motifs in *Taenopygia guttata* and *Xenopus laevis*. (D) ZNF410 orthologs binding motifs in *Danio rerio*. (E) Browser tracks of endogenous Zfp410 occupancy in the *Chd4* enhancer regions in mice G1E-ER4 cells. (F) The conserved sequence of ZNF410 binding motif in each ZNF410 peaks.

**Figure S5.**
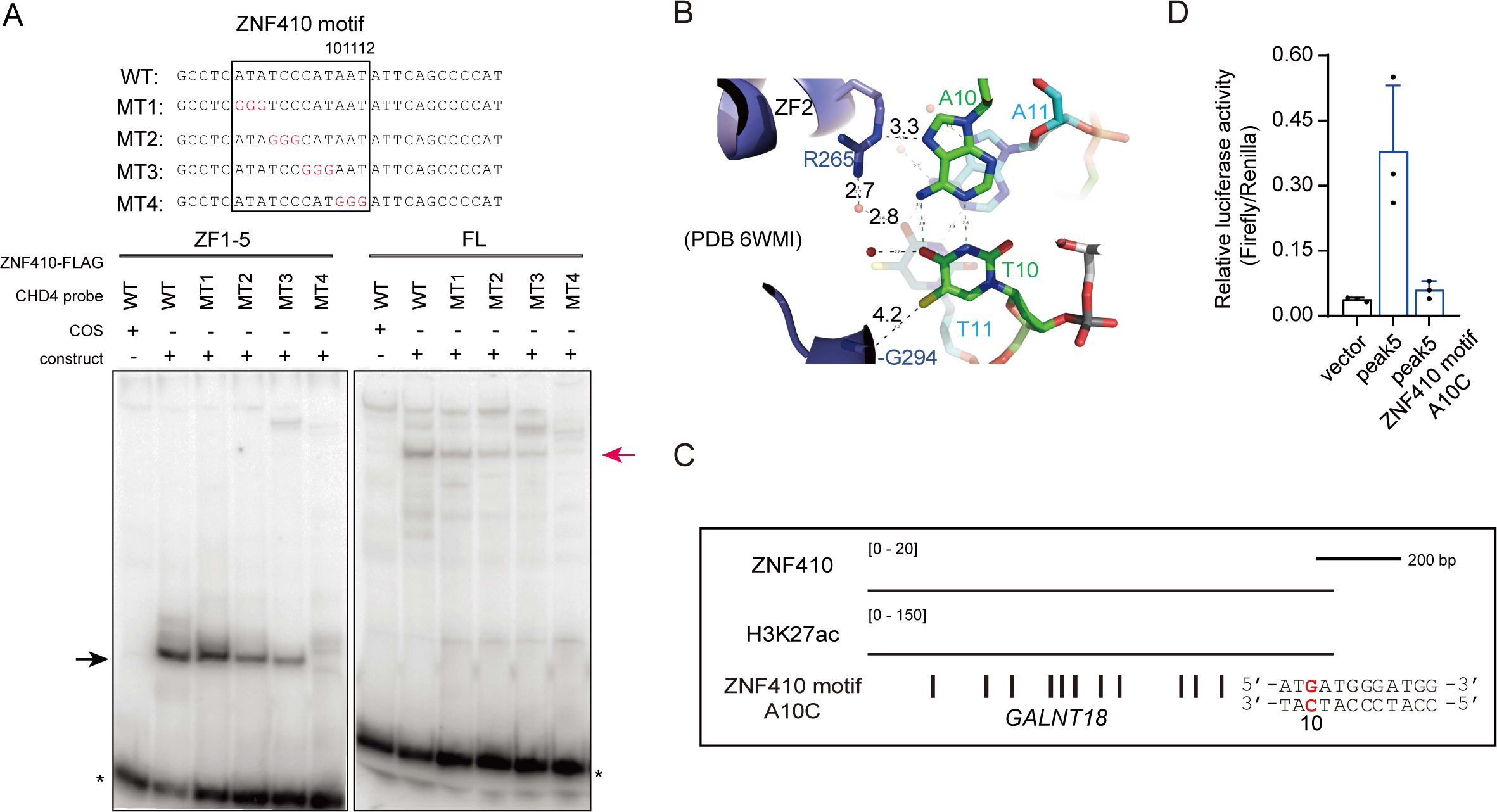
Identifying the key nucleotides of the ZNF410 motifs *in vitro* and *in vivo*, related to Figure 4. (A) Determination of the key nucleotide of ZNF410 binding motif via EMSA experiments. Probe sequences are shown above and mutant nucleotides are highlighted in red. A ‘COS emptý extract was used as negative control. Black arrow, ZF-probe complex; red arrow, ZNF410-probe complex; *, free probes; ZF, zinc finger; FL, full-length. (B) The interaction between ZF2 and A10 and A11 within ZNF410 binding motif. (C) ChIP-seq profiles of ZNF410 and H3K27ac occupation in the *GLANT18* locus. Homotypic mutated ZNF410 clustered motifs are shown below. (D) Results of luciferase reporter assays to measure the ZNF410 A10C mutated motif. Results are shown as mean ± SD (n=3).

**Figure S6.**
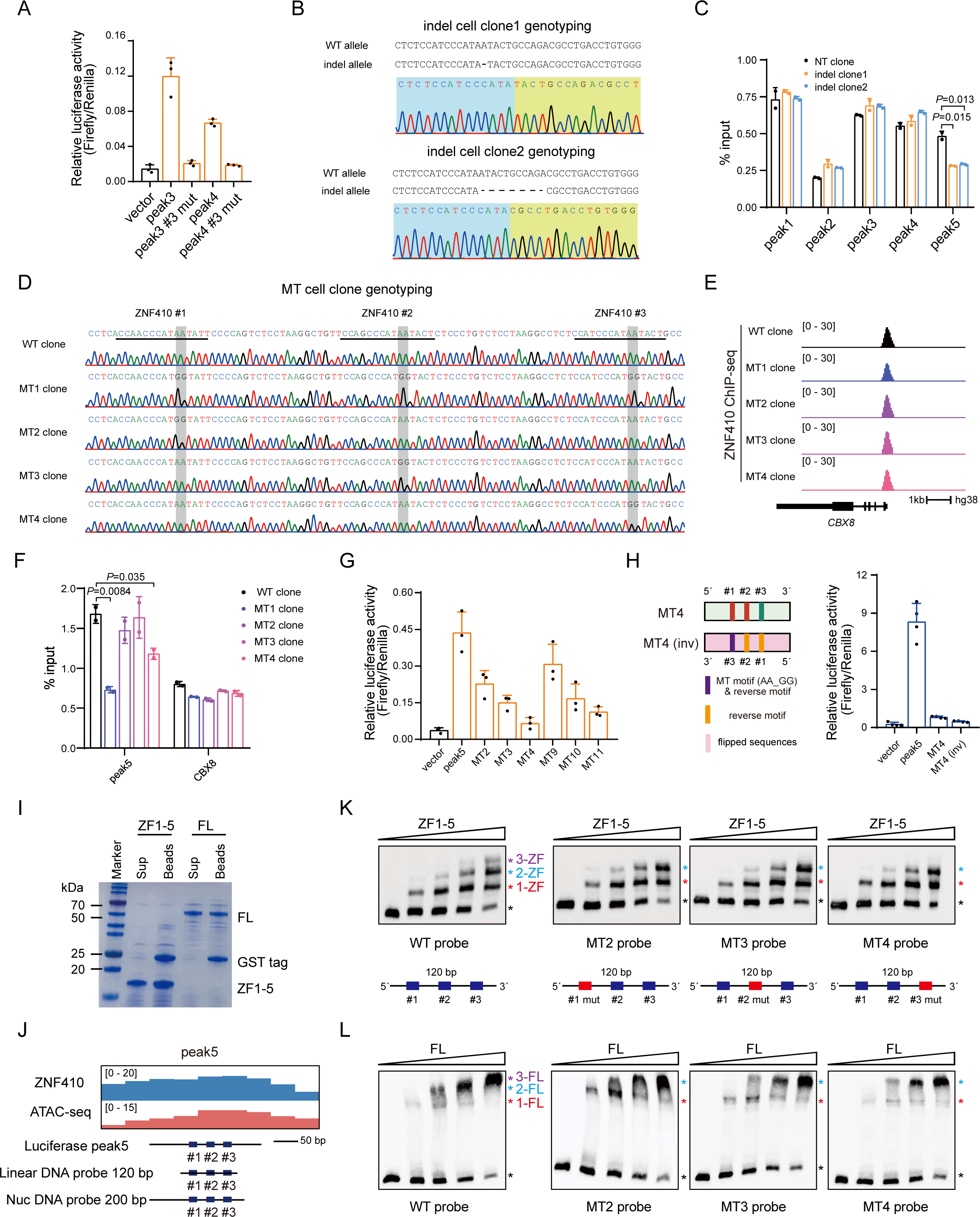
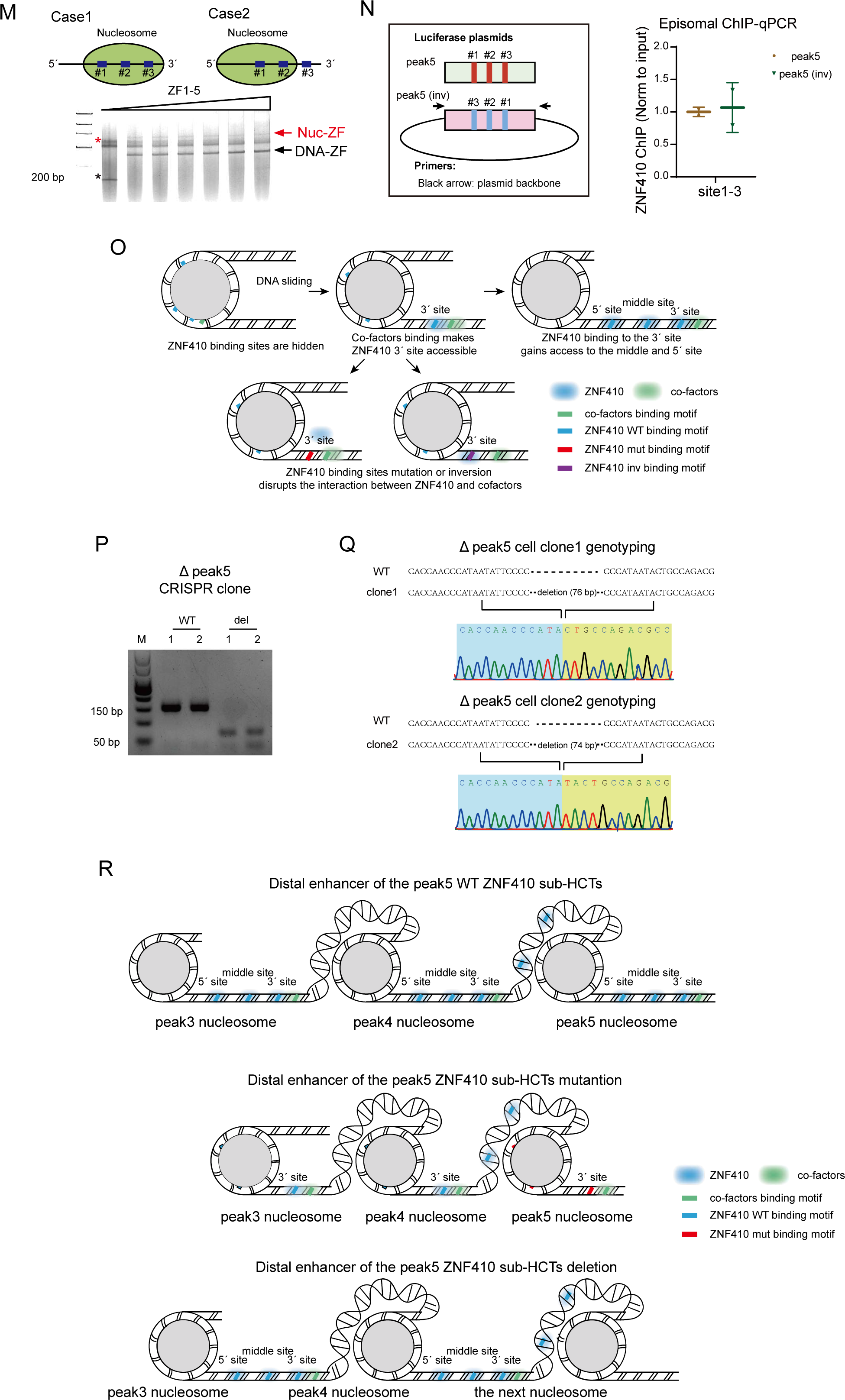
Site-dependent effect of the ZNF410 motifs *in vivo* and *in vitro*, related to Figure 5 and 6. (A) Luciferase assay results of peak3 and peak4 3’-ZNF410-motif mutated vectors. Results are shown as mean ± SD (n=3). #3, 3’ ZNF410 binding motif. (B) Sanger sequencing traces of the two peak5 3’-ZNF410-motif mutated indel single-cell clones. (C) ZNF410 ChIP-qPCR results of two peak5 3’-ZNF410-motif mutated indel single-cell clones in each ZNF410 peaks. Results are shown as mean ± SD (n=2). *P* values were calculated with unpaired Student’s *t*-test. (D) Sanger sequencing traces of each ZNF410 motif mutation homozygous single-cell clones. ZNF410 motifs are highlighted by grey box. MT, mutation. (E) ChIP-seq profiles of ZNF410 occupation in the *CBX8* locus. (F) Quantification of ZNF410 enrichment by ChIP-qPCR. Results are shown as mean ± SD (n=2). *P* values were calculated with unpaired Student’s *t*-test. (G) Verifying the site-dependent role of ZNF410 motifs in mutated and reverse orientation motif sequences. Results are shown as mean ± SD (n=3). (H) Luciferase assay results of inversion of whole MT4 sequences. Results are shown as mean ± SD (n=4). (I) The purified ZNF410 ZF1-5 and FL was subjected to SDS-PAGE (SDS–polyacrylamide gel electrophoresis) followed by Coomassie blue staining. Sup, supernatant. (J) Schematic of the location of linear DNA probe and nucleosome DNA probe. Nuc, nucleosome. (K-L) EMSA experiments reveal the formation of ZNF410 ZF1-5 (K) and FL (L) with DNA probe variants. Black asterisk, free probes; red asterisk, 1-ZF/FL-DNA complex; blue asterisk, 2-ZF/FL-DNA complex; purple asterisk, 3-ZF/FL-DNA complex; mut, mutation. (M) EMSA experiments reveal the formation of ZNF410 ZF1-5 with the nucleosome. Red arrow, ZF-nucleosome complex; black arrow, ZF-DNA complex. (N) Episomal ZNF410 ChIP-qPCR for K562 cells transfected with the peak5 and peak5 inv luciferase plasmids. Results are shown as mean ± SD (n=2). (O) A proposed model of ZNF410 cooperatively binding to sub-HCTs within the peak5. (P-Q) Genotyping of the homozygous peak5 knockouts single-cell CRISPR clones. (R) A proposed model of the ZNF410 binding to the distal enhancer. ZNF410 binding to the sub-HCTs within the peak5 gains access to the peak3 and peak4 of the whole distal enhancer.

**Table S1.**
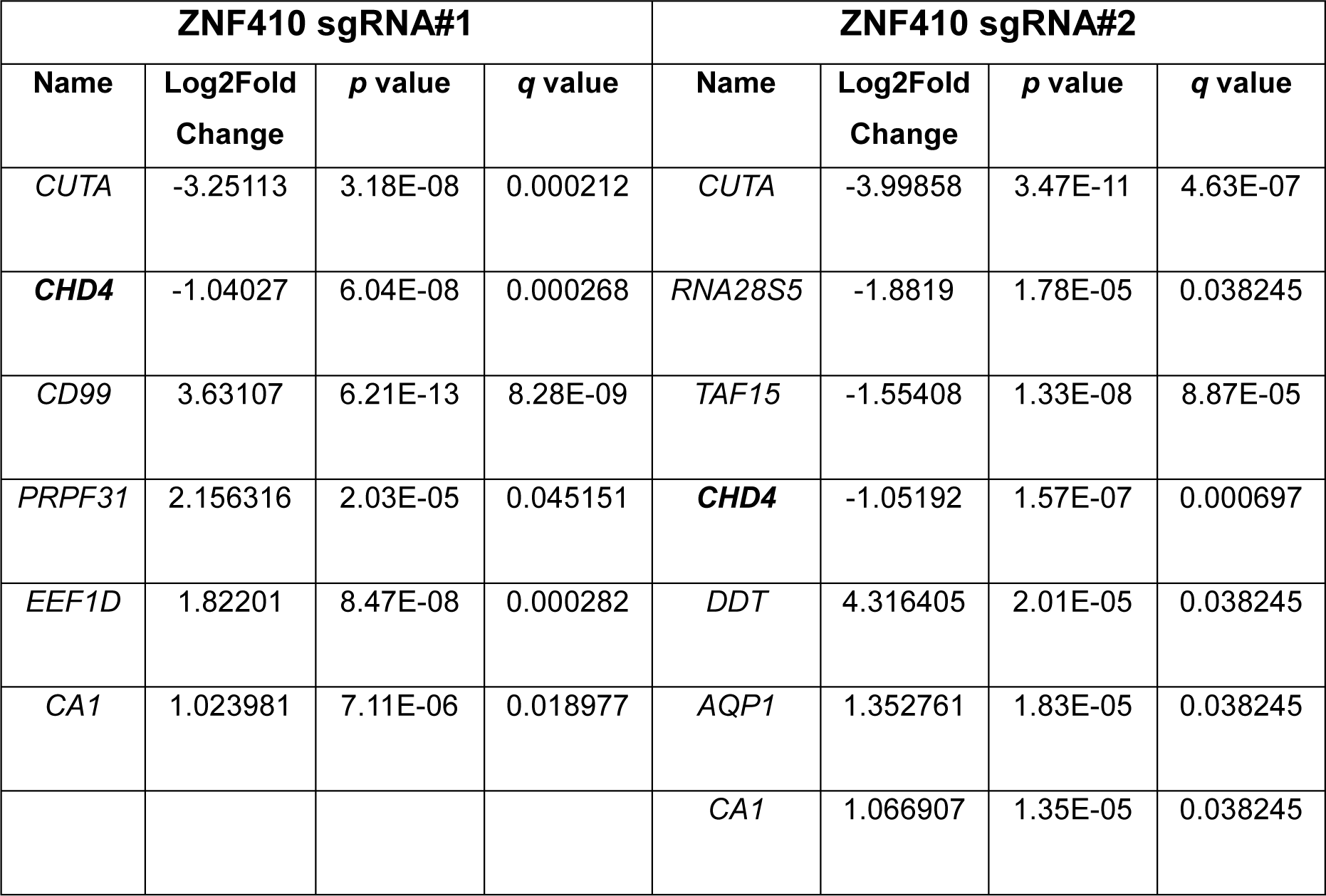
Differential gene analysis in K562 cells by RNA-seq, related to Figure 1.

| **ZNF410 sgRNA#1** |  |  |  | **ZNF410 sgRNA#2** |  |  |  |
| --- | --- | --- | --- | --- | --- | --- | --- |
| **Name** | **Log2Fold Change** | **p value** | **q value** | **Name** | **Log2Fold Change** | **p value** | **q value** |
| *CUTA* | -3.25113 | 3.18E-08 | 0.000212 | *CUTA* | -3.99858 | 3.47E-11 | 4.63E-07 |
| ***CHD4*** | -1.04027 | 6.04E-08 | 0.000268 | *RNA28S5* | -1.8819 | 1.78E-05 | 0.038245 |
| *CD99* | 3.63107 | 6.21E-13 | 8.28E-09 | *TAF15* | -1.55408 | 1.33E-08 | 8.87E-05 |
| *PRPF31* | 2.156316 | 2.03E-05 | 0.045151 | ***CHD4*** | -1.05192 | 1.57E-07 | 0.000697 |
| *EEF1D* | 1.82201 | 8.47E-08 | 0.000282 | *DDT* | 4.316405 | 2.01E-05 | 0.038245 |
| *CA1* | 1.023981 | 7.11E-06 | 0.018977 | *AQP1* | 1.352761 | 1.83E-05 | 0.038245 |
|  |  |  |  | *CA1* | 1.066907 | 1.35E-05 | 0.038245 |

**Table S2.**
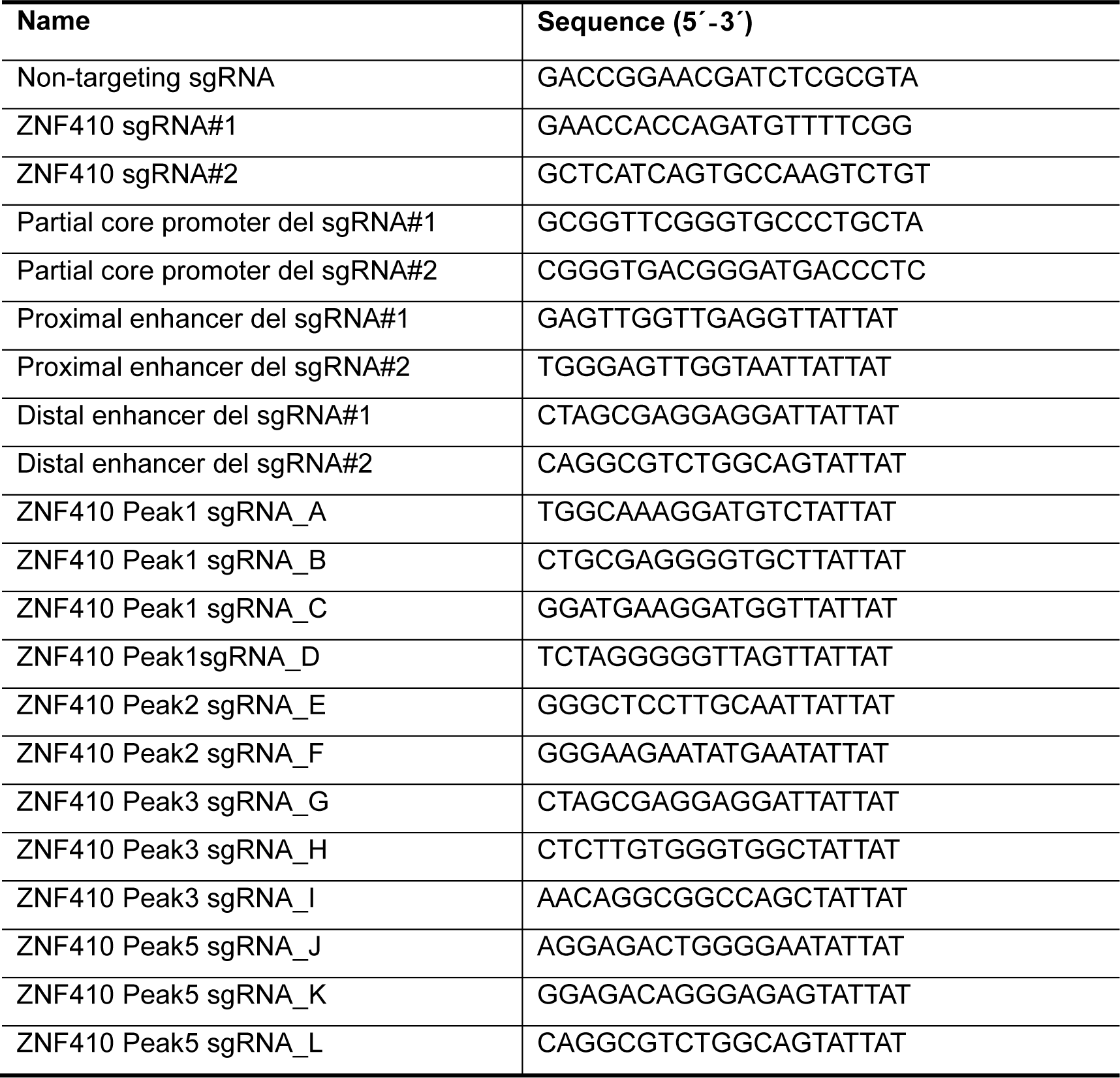
CRISPR guide RNA sequences, related to STAR METHODS.

**Table S3.**
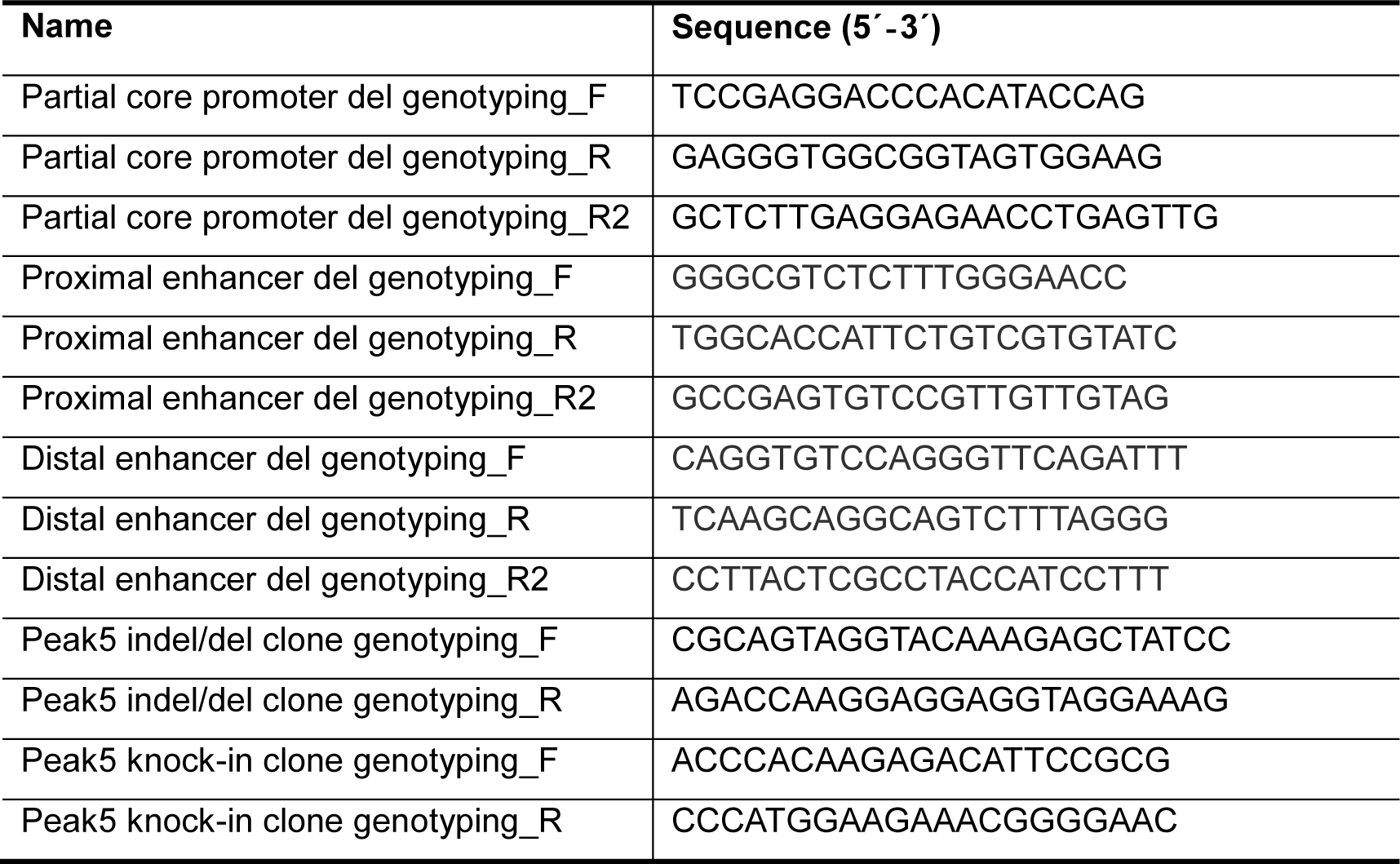
Primers for single cell genotyping, related to STAR METHODS.

**Table S4.**
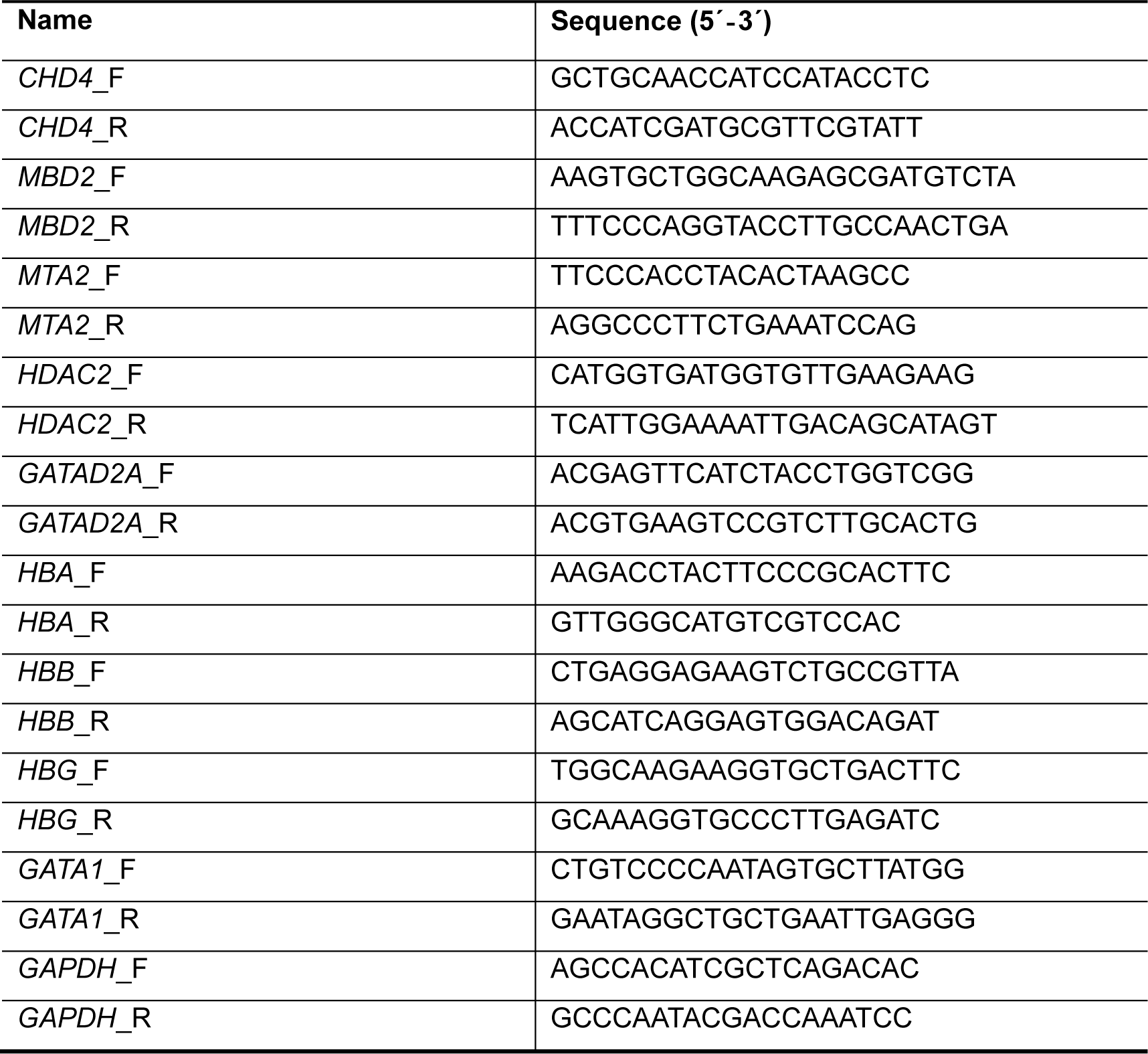
Primers for RT-qPCR, related to STAR METHODS.

**Table S5.**
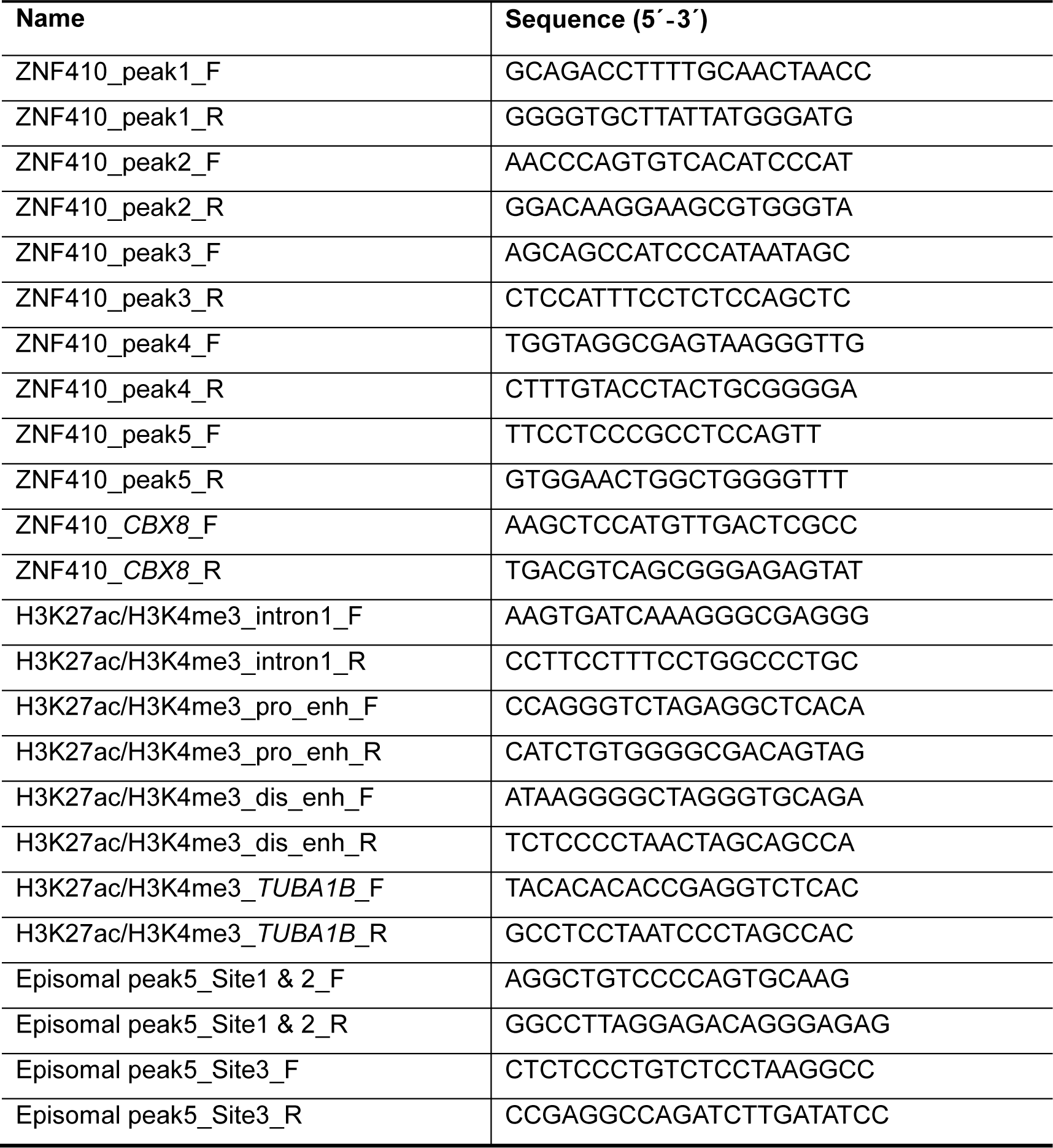
Primers for ChIP-qPCR, related to STAR METHODS.

## REFERENCES

1. Kim, S., and Wysocka, J. (2023). Deciphering the multi-scale, quantitative cis-regulatory code. Mol. Cell 83, 373–392. 10.1016/j.molcel.2022.12.032.

2. Banerji, J., Rusconi, S., and Schaffner, W. (1981). Expression of a β-globin gene is enhanced by remote SV40 DNA sequences. Cell 27, 299–308. 10.1016/0092-8674(81)90413-x.

3. Boyle, A.P., Davis, S., Shulha, H.P., Meltzer, P., Margulies, E.H., Weng, Z., Furey, T.S., and Crawford, G.E. (2008). High-resolution mapping and characterization of open chromatin across the genome. Cell 132, 311–322. 10.1016/j.cell.2007.12.014.

4. Kim, T.K., Hemberg, M., Gray, J.M., Costa, A.M., Bear, D.M., Wu, J., Harmin, D.A., Laptewicz, M., Barbara-Haley, K., Kuersten, S., et al. (2010). Widespread transcription at neuronal activity-regulated enhancers. Nature 465, 182–187. 10.1038/nature09033.

5. Visel, A., Blow, M.J., Li, Z., Zhang, T., Akiyama, J.A., Holt, A., Plajzer-Frick, I., Shoukry, M., Wright, C., Chen, F., et al. (2009). ChIP-seq accurately predicts tissue-specific activity of enhancers. Nature 457, 854–858. 10.1038/nature07730.

6. Heintzman, N.D., Hon, G.C., Hawkins, R.D., Kheradpour, P., Stark, A., Harp, L.F., Ye, Z., Lee, L.K., Stuart, R.K., Ching, C.W., et al. (2009). Histone modifications at human enhancers reflect global cell-type-specific gene expression. Nature 459, 108–112. 10.1038/nature07829.

7. Creyghton, M.P., Cheng, A.W., Welstead, G.G., Kooistra, T., Carey, B.W., Steine, E.J., Hanna, J., Lodato, M.A., Frampton, G.M., Sharp, P.A., et al. (2010). Histone H3K27ac separates active from poised enhancers and predicts developmental state. Proc. Natl. Acad. Sci. USA 107, 21931–21936. 10.1073/pnas.1016071107.

8. Andersson, R., Gebhard, C., Miguel-Escalada, I., Hoof, I., Bornholdt, J., Boyd, M., Chen, Y., Zhao, X., Schmidl, C., Suzuki, T., et al. (2014). An atlas of active enhancers across human cell types and tissues. Nature 507, 455–461. 10.1038/nature12787.

9. Bender, M.A., Ragoczy, T., Lee, J., Byron, R., Telling, A., Dean, A., and Groudine, M. (2012). The hypersensitive sites of the murine beta-globin locus control region act independently to affect nuclear localization and transcriptional elongation. Blood 119, 3820–3827. 10.1182/blood-2011-09-380485.

10. Hay, D., Hughes, J.R., Babbs, C., Davies, J.O.J., Graham, B.J., Hanssen, L., Kassouf, M.T., Marieke Oudelaar, A.M., Sharpe, J.A., Suciu, M.C., et al. (2016). Genetic dissection of the alpha-globin super-enhancer in vivo. Nat. Genet. 48, 895–903. 10.1038/ng.3605.

11. Will, A.J., Cova, G., Osterwalder, M., Chan, W.L., Wittler, L., Brieske, N., Heinrich, V., de Villartay, J.P., Vingron, M., Klopocki, E., et al. (2017). Composition and dosage of a multipartite enhancer cluster control developmental expression of Ihh (Indian hedgehog). Nat. Genet. 49, 1539–1545. 10.1038/ng.3939.

12. Frankel, N., Davis, G.K., Vargas, D., Wang, S., Payre, F., and Stern, D.L. (2010). Phenotypic robustness conferred by apparently redundant transcriptional enhancers. Nature 466, 490–493. 10.1038/nature09158.

13. Antosova, B., Smolikova, J., Klimova, L., Lachova, J., Bendova, M., Kozmikova, I., Machon, O., and Kozmik, Z. (2016). The Gene Regulatory Network of Lens Induction Is Wired through Meis-Dependent Shadow Enhancers of Pax6. PLoS Genet. 12, e1006441. 10.1371/journal.pgen.1006441.

14. Osterwalder, M., Barozzi, I., Tissieres, V., Fukuda-Yuzawa, Y., Mannion, B.J., Afzal, S.Y., Lee, E.A., Zhu, Y., Plajzer-Frick, I., Pickle, C.S., et al. (2018). Enhancer redundancy provides phenotypic robustness in mammalian development. Nature 554, 239–243. 10.1038/nature25461.

15. Shin, H.Y., Willi, M., HyunYoo, K., Zeng, X., Wang, C., Metser, G., and Hennighausen, L. (2016). Hierarchy within the mammary STAT5-driven Wap super-enhancer. Nat. Genet. 48, 904–911. 10.1038/ng.3606.

16. Huang, J., Li, K., Cai, W., Liu, X., Zhang, Y., Orkin, S.H., Xu, J., and Yuan, G.C. (2018). Dissecting super-enhancer hierarchy based on chromatin interactions. Nat. Commun. 9, 943. 10.1038/s41467-018-03279-9.

17. Perry, M.W., Boettiger, A.N., and Levine, M. (2011). Multiple enhancers ensure precision of gap gene-expression patterns in the Drosophila embryo. Proc. Natl. Acad. Sci. USA 108, 13570–13575. 10.1073/pnas.1109873108.

18. Lin, X., Liu, Y., Liu, S., Zhu, X., Wu, L., Zhu, Y., Zhao, D., Xu, X., Chemparathy, A., Wang, H., et al. (2022). Nested epistasis enhancer networks for robust genome regulation. Science 377, 1077–1085. 10.1126/science.abk3512.

19. Brosh, R., Coelho, C., Ribeiro-Dos-Santos, A.M., Ellis, G., Hogan, M.S., Ashe, H.J., Somogyi, N., Ordonez, R., Luther, R.D., Huang, E., et al. (2023). Synthetic regulatory genomics uncovers enhancer context dependence at the Sox2 locus. Mol. Cell 83, 1140–1152 e1147. 10.1016/j.molcel.2023.02.027.

20. Gotea, V., Visel, A., Westlund, J.M., Nobrega, M.A., Pennacchio, L.A., and Ovcharenko, I. (2010). Homotypic clusters of transcription factor binding sites are a key component of human promoters and enhancers. Genome Res. 20, 565–577. 10.1101/gr.104471.109.

21. Kato, G.J., Piel, F.B., Reid, C.D., Gaston, M.H., Ohene-Frempong, K., Krishnamurti, L., Smith, W.R., Panepinto, J.A., Weatherall, D.J., Costa, F.F., and Vichinsky, E.P. (2018). Sickle cell disease. Nat. Rev. Dis. Primers 4, 18010. 10.1038/nrdp.2018.10.

22. Taher, A.T., Musallam, K.M., and Cappellini, M.D. (2021). beta-Thalassemias. N. Engl. J. Med. 384, 727–743. 10.1056/NEJMra2021838.

23. Lan, X., Ren, R., Feng, R., Ly, L.C., Lan, Y., Zhang, Z., Aboreden, N., Qin, K., Horton, J.R., Grevet, J.D., et al. (2021). ZNF410 Uniquely Activates the NuRD Component CHD4 to Silence Fetal Hemoglobin Expression. Mol. Cell 81, 239–254 e238. 10.1016/j.molcel.2020.11.006.

24. Vinjamur, D.S., Yao, Q., Cole, M.A., McGuckin, C., Ren, C., Zeng, J., Hossain, M., Luk, K., Wolfe, S.A., Pinello, L., and Bauer, D.E. (2021). ZNF410 represses fetal globin by singular control of CHD4. Nat. Genet. 53, 719–728. 10.1038/s41588-021-00843-w.

25. Huang, P., Peslak, S.A., Ren, R., Khandros, E., Qin, K., Keller, C.A., Giardine, B., Bell, H.W., Lan, X., Sharma, M., et al. (2022). HIC2 controls developmental hemoglobin switching by repressing BCL11A transcription. Nat. Genet. 10.1038/s41588-022-01152-6.

26. Qin, K., Huang, P., Feng, R., Keller, C.A., Peslak, S.A., Khandros, E., Saari, M.S., Lan, X., Mayuranathan, T., Doerfler, P.A., et al. (2022). Dual function NFI factors control fetal hemoglobin silencing in adult erythroid cells. Nat. Genet. 10.1038/s41588-022-01076-1.

27. Amaya, M., Desai, M., Gnanapragasam, M.N., Wang, S.Z., Zu Zhu, S., Williams, D.C., Jr., and Ginder, G.D. (2013). Mi2beta-mediated silencing of the fetal gamma-globin gene in adult erythroid cells. Blood 121, 3493–3501. 10.1182/blood-2012-11-466227.

28. Kaur, G., Ren, R., Hammel, M., Horton, J.R., Yang, J., Cao, Y., He, C., Lan, F., Lan, X., Blobel, G.A., et al. (2023). Allosteric autoregulation of DNA binding via a DNA-mimicking protein domain: a biophysical study of ZNF410-DNA interaction using small angle X-ray scattering. Nucleic Acids Res. 51, 1674–1686. 10.1093/nar/gkac1274.

29. Fagerberg, L., Hallstrom, B.M., Oksvold, P., Kampf, C., Djureinovic, D., Odeberg, J., Habuka, M., Tahmasebpoor, S., Danielsson, A., Edlund, K., et al. (2014). Analysis of the human tissue-specific expression by genome-wide integration of transcriptomics and antibody-based proteomics. Mol. Cell Proteomics 13, 397–406. 10.1074/mcp.M113.035600.

30. Tabula Sapiens, C., Jones, R.C., Karkanias, J., Krasnow, M.A., Pisco, A.O., Quake, S.R., Salzman, J., Yosef, N., Bulthaup, B., Brown, P., et al. (2022). The Tabula Sapiens: A multiple-organ, single-cell transcriptomic atlas of humans. Science 376, eabl4896. 10.1126/science.abl4896.

31. King, A.J., Songdej, D., Downes, D.J., Beagrie, R.A., Liu, S., Buckley, M., Hua, P., Suciu, M.C., Marieke Oudelaar, A., Hanssen, L.L.P., et al. (2021). Reactivation of a developmentally silenced embryonic globin gene. Nat. Commun. 12, 4439. 10.1038/s41467-021-24402-3.

32. Consortium, E.P., Moore, J.E., Purcaro, M.J., Pratt, H.E., Epstein, C.B., Shoresh, N., Adrian, J., Kawli, T., Davis, C.A., Dobin, A., et al. (2020). Expanded encyclopaedias of DNA elements in the human and mouse genomes. Nature 583, 699–710. 10.1038/s41586-020-2493-4.

33. Carleton, J.B., Berrett, K.C., and Gertz, J. (2017). Multiplex Enhancer Interference Reveals Collaborative Control of Gene Regulation by Estrogen Receptor alpha-Bound Enhancers. Cell Syst. 5, 333–344 e335. 10.1016/j.cels.2017.08.011.

34. Kvon, E.Z., Waymack, R., Gad, M., and Wunderlich, Z. (2021). Enhancer redundancy in development and disease. Nat. Rev. Genet. 22, 324–336. 10.1038/s41576-020-00311-x.

35. Hoffman, J.A., Trotter, K.W., Day, C.R., Ward, J.M., Inoue, K., Rodriguez, J., and Archer, T.K. (2022). Multimodal regulatory elements within a hormone-specific super enhancer control a heterogeneous transcriptional response. Mol. Cell 82, 803–815 e805. 10.1016/j.molcel.2021.12.035.

36. O’Shaughnessy-Kirwan, A., Signolet, J., Costello, I., Gharbi, S., and Hendrich, B. (2015). Constraint of gene expression by the chromatin remodelling protein CHD4 facilitates lineage specification. Development 142, 2586–2597. 10.1242/dev.125450.

37. Gilbert, L.A., Larson, M.H., Morsut, L., Liu, Z., Brar, G.A., Torres, S.E., Stern-Ginossar, N., Brandman, O., Whitehead, E.H., Doudna, J.A., et al. (2013). CRISPR-mediated modular RNA-guided regulation of transcription in eukaryotes. Cell 154, 442–451. 10.1016/j.cell.2013.06.044.

38. Schultz, D.C., Ayyanathan, K., Negorev, D., Maul, G.G., and Rauscher, F.J., 3rd (2002). SETDB1: a novel KAP-1-associated histone H3, lysine 9-specific methyltransferase that contributes to HP1-mediated silencing of euchromatic genes by KRAB zinc-finger proteins. Genes Dev. 16, 919–932. 10.1101/gad.973302.

39. Groner, A.C., Meylan, S., Ciuffi, A., Zangger, N., Ambrosini, G., Denervaud, N., Bucher, P., and Trono, D. (2010). KRAB-zinc finger proteins and KAP1 can mediate long-range transcriptional repression through heterochromatin spreading. PLoS Genet. 6, e1000869. 10.1371/journal.pgen.1000869.

40. Liu, N., Hargreaves, V.V., Zhu, Q., Kurland, J.V., Hong, J., Kim, W., Sher, F., Macias-Trevino, C., Rogers, J.M., Kurita, R., et al. (2018). Direct Promoter Repression by BCL11A Controls the Fetal to Adult Hemoglobin Switch. Cell 173, 430–442 e417. 10.1016/j.cell.2018.03.016.

41. Long, H.K., Osterwalder, M., Welsh, I.C., Hansen, K., Davies, J.O.J., Liu, Y.E., Koska, M., Adams, A.T., Aho, R., Arora, N., et al. (2020). Loss of Extreme Long-Range Enhancers in Human Neural Crest Drives a Craniofacial Disorder. Cell Stem Cell 27, 765–783 e714. 10.1016/j.stem.2020.09.001.

42. Ezer, D., Zabet, N.R., and Adryan, B. (2014). Homotypic clusters of transcription factor binding sites: A model system for understanding the physical mechanics of gene expression. Comput. Struct. Biotechnol. J. 10, 63–69. 10.1016/j.csbj.2014.07.005.

43. Gordan, R., Shen, N., Dror, I., Zhou, T., Horton, J., Rohs, R., and Bulyk, M.L. (2013). Genomic regions flanking E-box binding sites influence DNA binding specificity of bHLH transcription factors through DNA shape. Cell Rep. 3, 1093–1104. 10.1016/j.celrep.2013.03.014.

44. Yella, V.R., Bhimsaria, D., Ghoshdastidar, D., Rodriguez-Martinez, J.A., Ansari, A.Z., and Bansal, M. (2018). Flexibility and structure of flanking DNA impact transcription factor affinity for its core motif. Nucleic Acids Res. 46, 11883–11897. 10.1093/nar/gky1057.

45. Adams, C.C., and Workman, J.L. (1995). Binding of disparate transcriptional activators to nucleosomal DNA is inherently cooperative. Mol. Cell Biol. 15, 1405–1421. 10.1128/MCB.15.3.1405.

46. Polach, K.J., and Widom, J. (1996). A model for the cooperative binding of eukaryotic regulatory proteins to nucleosomal target sites. J. Mol. Biol. 258, 800–812. 10.1006/jmbi.1996.0288.

47. Vashee, S., Melcher, K., Ding, W.V., Johnston, S.A., and Kodadek, T. (1998). Evidence for two modes of cooperative DNA binding in vivo that do not involve direct protein-protein interactions. Curr. Biol. 8, 452–458. 10.1016/s0960-9822(98)70179-4.

48. Mirny, L.A. (2010). Nucleosome-mediated cooperativity between transcription factors. Proc. Natl. Acad. Sci. USA 107, 22534–22539. 10.1073/pnas.0913805107.

49. Suganuma, T., Gutierrez, J.L., Li, B., Florens, L., Swanson, S.K., Washburn, M.P., Abmayr, S.M., and Workman, J.L. (2008). ATAC is a double histone acetyltransferase complex that stimulates nucleosome sliding. Nat. Struct. Mol. Biol. 15, 364–372. 10.1038/nsmb.1397.

50. Zaret, K.S., and Carroll, J.S. (2011). Pioneer transcription factors: establishing competence for gene expression. Genes Dev. 25, 2227–2241. 10.1101/gad.176826.111.

51. Zhu, F., Farnung, L., Kaasinen, E., Sahu, B., Yin, Y., Wei, B., Dodonova, S.O., Nitta, K.R., Morgunova, E., Taipale, M., et al. (2018). The interaction landscape between transcription factors and the nucleosome. Nature 562, 76–81. 10.1038/s41586-018-0549-5.

52. Dominguez, A.A., Lim, W.A., and Qi, L.S. (2016). Beyond editing: repurposing CRISPR-Cas9 for precision genome regulation and interrogation. Nat. Rev. Mol. Cell Bio. 17, 5–15. 10.1038/nrm.2015.2.

53. Lambert, S.A., Jolma, A., Campitelli, L.F., Das, P.K., Yin, Y., Albu, M., Chen, X., Taipale, J., Hughes, T.R., and Weirauch, M.T. (2018). The Human Transcription Factors. Cell 172, 650–665. 10.1016/j.cell.2018.01.029.

54. Giorgetti, L., Siggers, T., Tiana, G., Caprara, G., Notarbartolo, S., Corona, T., Pasparakis, M., Milani, P., Bulyk, M.L., and Natoli, G. (2010). Noncooperative interactions between transcription factors and clustered DNA binding sites enable graded transcriptional responses to environmental inputs. Mol. Cell 37, 418–428. 10.1016/j.molcel.2010.01.016.

55. Balestrieri, C., Alfarano, G., Milan, M., Tosi, V., Prosperini, E., Nicoli, P., Palamidessi, A., Scita, G., Diaferia, G.R., and Natoli, G. (2018). Co-optation of Tandem DNA Repeats for the Maintenance of Mesenchymal Identity. Cell 173, 1150–1164 e1114. 10.1016/j.cell.2018.03.081.

56. Johnson, A.D., Meyer, B.J., and Ptashne, M. (1979). Interactions between DNA-bound repressors govern regulation by the lambda phage repressor. Proc. Natl. Acad. Sci. USA 76, 5061–5065. 10.1073/pnas.76.10.5061.

57. Stayrook, S., Jaru-Ampornpan, P., Ni, J., Hochschild, A., and Lewis, M. (2008). Crystal structure of the lambda repressor and a model for pairwise cooperative operator binding. Nature 452, 1022–1025. 10.1038/nature06831.

58. Burz, D.S., Rivera-Pomar, R., Jackle, H., and Hanes, S.D. (1998). Cooperative DNA-binding by Bicoid provides a mechanism for threshold-dependent gene activation in the Drosophila embryo. EMBO J. 17, 5998–6009. 10.1093/emboj/17.20.5998.

59. Lebrecht, D., Foehr, M., Smith, E., Lopes, F.J., Vanario-Alonso, C.E., Reinitz, J., Burz, D.S., and Hanes, S.D. (2005). Bicoid cooperative DNA binding is critical for embryonic patterning in Drosophila. Proc. Natl. Acad. Sci. USA 102, 13176–13181. 10.1073/pnas.0506462102.

60. Smith, R.P., Taher, L., Patwardhan, R.P., Kim, M.J., Inoue, F., Shendure, J., Ovcharenko, I., and Ahituv,N. (2013). Massively parallel decoding of mammalian regulatory sequences supports a flexible organizational model. Nat. Genet. 45, 1021–1028. 10.1038/ng.2713.

61. Grossman, S.R., Zhang, X., Wang, L., Engreitz, J., Melnikov, A., Rogov, P., Tewhey, R., Isakova, A., Deplancke, B., Bernstein, B.E., et al. (2017). Systematic dissection of genomic features determining transcription factor binding and enhancer function. Proc. Natl. Acad. Sci. USA 114, E1291–E1300. 10.1073/pnas.1621150114.

62. Georgakopoulos-Soares, I., Deng, C., Agarwal, V., Chan, C.S.Y., Zhao, J., Inoue, F., and Ahituv, N. (2023). Transcription factor binding site orientation and order are major drivers of gene regulatory activity. Nat. Commun. 14, 2333. 10.1038/s41467-023-37960-5.

63. Soufi, A., Garcia, M.F., Jaroszewicz, A., Osman, N., Pellegrini, M., and Zaret, K.S. (2015). Pioneer transcription factors target partial DNA motifs on nucleosomes to initiate reprogramming. Cell 161, 555–568. 10.1016/j.cell.2015.03.017.

64. Sinha, K.K., Bilokapic, S., Du, Y., Malik, D., and Halic, M. (2023). Histone modifications regulate pioneer transcription factor cooperativity. Nature 619, 378–384. 10.1038/s41586-023-06112-6.

65. Inukai, S., Kock, K.H., and Bulyk, M.L. (2017). Transcription factor-DNA binding: beyond binding site motifs. Curr. Opin. Genet. Dev. 43, 110–119. 10.1016/j.gde.2017.02.007.

66. Morgunova, E., and Taipale, J. (2017). Structural perspective of cooperative transcription factor binding. Curr. Opin. Struct. Biol. 47, 1–8. 10.1016/j.sbi.2017.03.006.

67. Garvie, C.W., Hagman, J., and Wolberger, C. (2001). Structural studies of Ets-1/Pax5 complex formation on DNA. Mol. Cell 8, 1267–1276. 10.1016/s1097-2765(01)00410-5.

68. Slattery, M., Riley, T., Liu, P., Abe, N., Gomez-Alcala, P., Dror, I., Zhou, T., Rohs, R., Honig, B., Bussemaker, H.J., and Mann, R.S. (2011). Cofactor binding evokes latent differences in DNA binding specificity between Hox proteins. Cell 147, 1270–1282. 10.1016/j.cell.2011.10.053.

69. Lelli, K.M., Slattery, M., and Mann, R.S. (2012). Disentangling the many layers of eukaryotic transcriptional regulation. Annu. Rev. Genet. 46, 43–68. 10.1146/annurev-genet-110711-155437.

70. Siggers, T., and Gordan, R. (2014). Protein-DNA binding: complexities and multi-protein codes. Nucleic Acids Res. 42, 2099–2111. 10.1093/nar/gkt1112.

71. Grubert, F., Zaugg, J.B., Kasowski, M., Ursu, O., Spacek, D.V., Martin, A.R., Greenside, P., Srivas, R., Phanstiel, D.H., Pekowska, A., et al. (2015). Genetic Control of Chromatin States in Humans Involves Local and Distal Chromosomal Interactions. Cell 162, 1051–1065. 10.1016/j.cell.2015.07.048.

72. Waszak, S.M., Delaneau, O., Gschwind, A.R., Kilpinen, H., Raghav, S.K., Witwicki, R.M., Orioli, A., Wiederkehr, M., Panousis, N.I., Yurovsky, A., et al. (2015). Population Variation and Genetic Control of Modular Chromatin Architecture in Humans. Cell 162, 1039–1050. 10.1016/j.cell.2015.08.001.

73. Kumasaka, N., Knights, A.J., and Gaffney, D.J. (2016). Fine-mapping cellular QTLs with RASQUAL and ATAC-seq. Nat. Genet. 48, 206–213. 10.1038/ng.3467.

74. Alasoo, K., Rodrigues, J., Mukhopadhyay, S., Knights, A.J., Mann, A.L., Kundu, K., Consortium, H., Hale, C., Dougan, G., and Gaffney, D.J. (2018). Shared genetic effects on chromatin and gene expression indicate a role for enhancer priming in immune response. Nat. Genet. 50, 424–431. 10.1038/s41588-018-0046-7.

75. Gate, R.E., Cheng, C.S., Aiden, A.P., Siba, A., Tabaka, M., Lituiev, D., Machol, I., Gordon, M.G., Subramaniam, M., Shamim, M., et al. (2018). Genetic determinants of co-accessible chromatin regions in activated T cells across humans. Nat. Genet. 50, 1140–1150. 10.1038/s41588-018-0156-2.

76. Kurita, R., Suda, N., Sudo, K., Miharada, K., Hiroyama, T., Miyoshi, H., Tani, K., and Nakamura, Y. (2013). Establishment of immortalized human erythroid progenitor cell lines able to produce enucleated red blood cells. PLoS One 8, e59890. 10.1371/journal.pone.0059890.

77. Patel, A., Hashimoto, H., Zhang, X., and Cheng, X. (2016). Characterization of How DNA Modifications Affect DNA Binding by C2H2 Zinc Finger Proteins. Methods Enzymol. 573, 387–401. 10.1016/bs.mie.2016.01.019.

78. Andrews, N.C., and Faller, D.V. (1991). A rapid micropreparation technique for extraction of DNA-binding proteins from limiting numbers of mammalian cells. Nucleic Acids Res. 19, 2499. 10.1093/nar/19.9.2499.

79. Luger, K., Rechsteiner, T.J., and Richmond, T.J. (1999). Expression and purification of recombinant histones and nucleosome reconstitution. Methods Mol. Biol. 119, 1–16. 10.1385/1-59259-681-9:1.

80. Corces, M.R., Trevino, A.E., Hamilton, E.G., Greenside, P.G., Sinnott-Armstrong, N.A., Vesuna, S., Satpathy, A.T., Rubin, A.J., Montine, K.S., Wu, B., et al. (2017). An improved ATAC-seq protocol reduces background and enables interrogation of frozen tissues. Nat. Methods 14, 959–962. 10.1038/nmeth.4396.

81. Kim, D., Paggi, J.M., Park, C., Bennett, C., and Salzberg, S.L. (2019). Graph-based genome alignment and genotyping with HISAT2 and HISAT-genotype. Nat. Biotechnol. 37, 907–915. 10.1038/s41587-019-0201-4.

82. Li, H., Handsaker, B., Wysoker, A., Fennell, T., Ruan, J., Homer, N., Marth, G., Abecasis, G., Durbin, R., and Genome Project Data Processing, S. (2009). The Sequence Alignment/Map format and SAMtools. Bioinformatics 25, 2078–2079. 10.1093/bioinformatics/btp352.

83. Anders, S., Pyl, P.T., and Huber, W. (2015). HTSeq--a Python framework to work with high-throughput sequencing data. Bioinformatics 31, 166–169. 10.1093/bioinformatics/btu638.

84. Li, B., and Dewey, C.N. (2011). RSEM: accurate transcript quantification from RNA-Seq data with or without a reference genome. BMC Bioinformatics 12, 323. 10.1186/1471-2105-12-323.

85. Love, M.I., Huber, W., and Anders, S. (2014). Moderated estimation of fold change and dispersion for RNA-seq data with DESeq2. Genome Biol. 15, 550. 10.1186/s13059-014-0550-8.

86. Langmead, B., and Salzberg, S.L. (2012). Fast gapped-read alignment with Bowtie 2. Nat. Methods 9, 357–359. 10.1038/nmeth.1923.

87. Ramirez, F., Ryan, D.P., Gruning, B., Bhardwaj, V., Kilpert, F., Richter, A.S., Heyne, S., Dundar, F., and Manke, T. (2016). deepTools2: a next generation web server for deep-sequencing data analysis. Nucleic Acids Res. 44, W160–165. 10.1093/nar/gkw257.

88. Zhang, Y., Liu, T., Meyer, C.A., Eeckhoute, J., Johnson, D.S., Bernstein, B.E., Nusbaum, C., Myers, R.M., Brown, M., Li, W., and Liu, X.S. (2008). Model-based analysis of ChIP-Seq (MACS). Genome Biol. 9, R137. 10.1186/gb-2008-9-9-r137.

89. Quinlan, A.R., and Hall, I.M. (2010). BEDTools: a flexible suite of utilities for comparing genomic features. Bioinformatics 26, 841–842. 10.1093/bioinformatics/btq033.

90. Heinz, S., Benner, C., Spann, N., Bertolino, E., Lin, Y.C., Laslo, P., Cheng, J.X., Murre, C., Singh, H., and Glass, C.K. (2010). Simple combinations of lineage-determining transcription factors prime cis-regulatory elements required for macrophage and B cell identities. Mol. Cell 38, 576–589. 10.1016/j.molcel.2010.05.004.

